# Sustained Relief of Chronic Pain via a Nav1.7-Targeting ASO–siRNA Conjugate

**DOI:** 10.64898/2026.03.27.714734

**Authors:** Bin Ren, Chang Yu, Weiqin Yin, Zhuangchuan Yuan, Hongyu Chen, Yicao Liu, Bin Fang, Suyuan Liu, Lianchao Gao, Zhiwei Cao, Qingqing Yu, Xin Qiu, Peixia Yu, Guohao Wang

## Abstract

Chronic pain affects billions globally, yet safe, long-lasting, and non-addictive analgesics remain lacking. Nav1.7 is a genetically validated pain target, but traditional small molecules have repeatedly failed. Therapeutic oligonucleotides-antisense oligonucleotides (ASOs) and siRNAs-offer selective, durable silencing. We developed N02C0702, an ASO-siRNA conjugate (ASC), achieving robust Nav1.7 knockdown and sustained analgesia without additional delivery vehicles. N02C0702 outperformed individual ASO (N02A114) and siRNA (N02S154) moieties at mRNA and protein levels and in pain relief. In CFA-induced inflammatory pain, a single intrathecal dose exceeded naproxen and suzetrigine, while in SNL neuropathic pain, efficacy persisted up to 56 days, comparable to or surpassing pregabalin. Genome-wide RNA sequencing confirmed minimal off-target effects. N02C0702 highlights Nav1.7 as a key analgesic target and demonstrates the ASC platform’s potential for chronic pain and other CNS-related pathologies, offering durable, selective, and safe therapeutic effects.

## INTRODUCTION

Chronic pain is a pervasive global health challenge, affecting billions and imposing profound physical, psychological, and economic burdens^1–3^. Current therapies are inadequate: opioids carry high addiction risk^4^ and are no longer first-line for most chronic pain syndromes^5,6^, while non-opioid alternatives, including NSAIDs, provide limited relief and are associated with gastrointestinal, cardiovascular, and renal complications^7^. Recently approved sodium channel–targeting analgesics, such as suzetrigine (VX-548), represent an important advance^8–11^, yet clinical trials in lumbosacral radiculopathy demonstrated efficacy comparable to placebo^12^. Moreover, existing analgesics often require frequent dosing and lack durable effects, underscoring the need for therapeutics that are effective, selective, and long-lasting.

Voltage-gated sodium (Nav) channels are central mediators of nociception. Of the nine subtypes (Nav1.1-1.9), Nav1.7, Nav1.8, and Nav1.9 play critical roles in pain signaling^13–15^. Nav1.7 acts as a threshold channel for action potential initiation and is tightly linked to human pain: loss-of-function mutations confer congenital insensitivity to pain^16–18^, whereas gain-of-function mutations drive inherited erythromelalgia, paroxysmal extreme pain disorder, and idiopathic small fiber neuropathy^19–22^. Despite intensive efforts, small molecules, peptides, and antibodies targeting Nav1.7 have largely failed clinically^23–29^. Challenges include high sequence homology among Nav isoforms, complicating selective inhibition^23,25,30^; off-target activity on Nav1.4 and Nav1.5 risks severe skeletal or cardiac toxicity^29,31^; insufficient engagement of both peripheral and central neurons^14^; and uncertainty regarding the extent of Nav1.7 blockade required for meaningful analgesia^16,32,33^.

Oligonucleotide therapeutics-including antisense oligonucleotides (ASOs) and small interfering RNAs (siRNAs)-offer unique solutions^34–41^. Sequence-specific base pairing enables selective silencing of individual isoforms and even allele-specific targeting^42–47^. Chemical modifications have enhanced stability, potency, and duration^48–50^, with siRNAs achieving exceptionally long-lasting effects, exemplified by Inclisiran’s six-month dosing schedule^51,52^. These properties make oligonucleotides well suited for chronic pain, where durable, subtype-specific inhibition is essential.

Here, we report a novel oligonucleotide scaffold, ASC (ASO–siRNA Conjugate), in which an ASO and an siRNA are covalently linked and both target Nav1.7. The lead compound, N02C0702, achieved robust and durable knockdown of Nav1.7 mRNA and protein in vitro and in vivo, without additional delivery vehicles. In rats with complete Freund’s adjuvant–induced inflammatory pain, N02C0702 outperformed suzetrigine and naproxen. In a spinal nerve ligation model of neuropathic pain, it exceeded the efficacy of its individual ASO or siRNA components and suzetrigine, while matching pregabalin. Remarkably, a single intrathecal dose provided analgesia for up to 56 days, whereas pregabalin and suzetrigine require multiple daily administrations^53,54^. These findings highlight the potential of oligonucleotide-based Nav1.7 targeting to provide potent, selective, and long-lasting analgesia, supporting the translation of ASCs as a new therapeutic modality for chronic pain.

## RESULTS

### In Vitro Screening of Nav1.7-Targeting ASOs and siRNAs

Therapeutic ASOs and siRNAs act via distinct mechanisms: ASOs induce RNase H–mediated degradation of target mRNA, whereas siRNAs silence genes through RNA interference (RNAi) via the RNA-induced silencing complex (RISC)^35,55^. To identify potent candidates, 129 oligonucleotides—46 ASOs and 83 siRNAs—were designed to target conserved regions of human and rat SCN9A transcripts encoding Nav1.7. ASOs were single-stranded gapmers with five 2’-O-methoxyethyl (MOE)-modified nucleotides flanking each end, while siRNAs were asymmetric duplexes with enhanced stabilization chemistry (ESC) modifications, including a 5’ (E)-vinylphosphonate (E-VP) on the guide strand (**Table S1, S2**).

High-throughput cell-based screening in human SK-N-AS cells using RT–qPCR ranked candidates by knockdown efficiency. ASOs reduced Nav1.7 mRNA by 21–86% at 10 nM and 51–97% at 100 nM (**Figure 1A**), whereas siRNAs achieved 6–80% and 19–88% reductions at 0.1 nM and 10 nM, respectively (**Figure 1B**). Considering that ∼50% knockdown may be insufficient for analgesia, >80% reduction (100 nM for ASOs, 10 nM for siRNAs) was used as a cutoff, yielding 28 ASOs and 12 siRNAs for further validation. Using primary rat DRG neurons as a physiologically relevant model, 10 of 28 ASOs were excluded due to <60% inhibition at 1 μM, while the top 12 siRNAs exhibited nanomolar IC₅₀ values (**Figure S1A-M**), resulting in 18 ASOs and 12 siRNAs advanced for in vitro safety testing.

**Figure 1.**
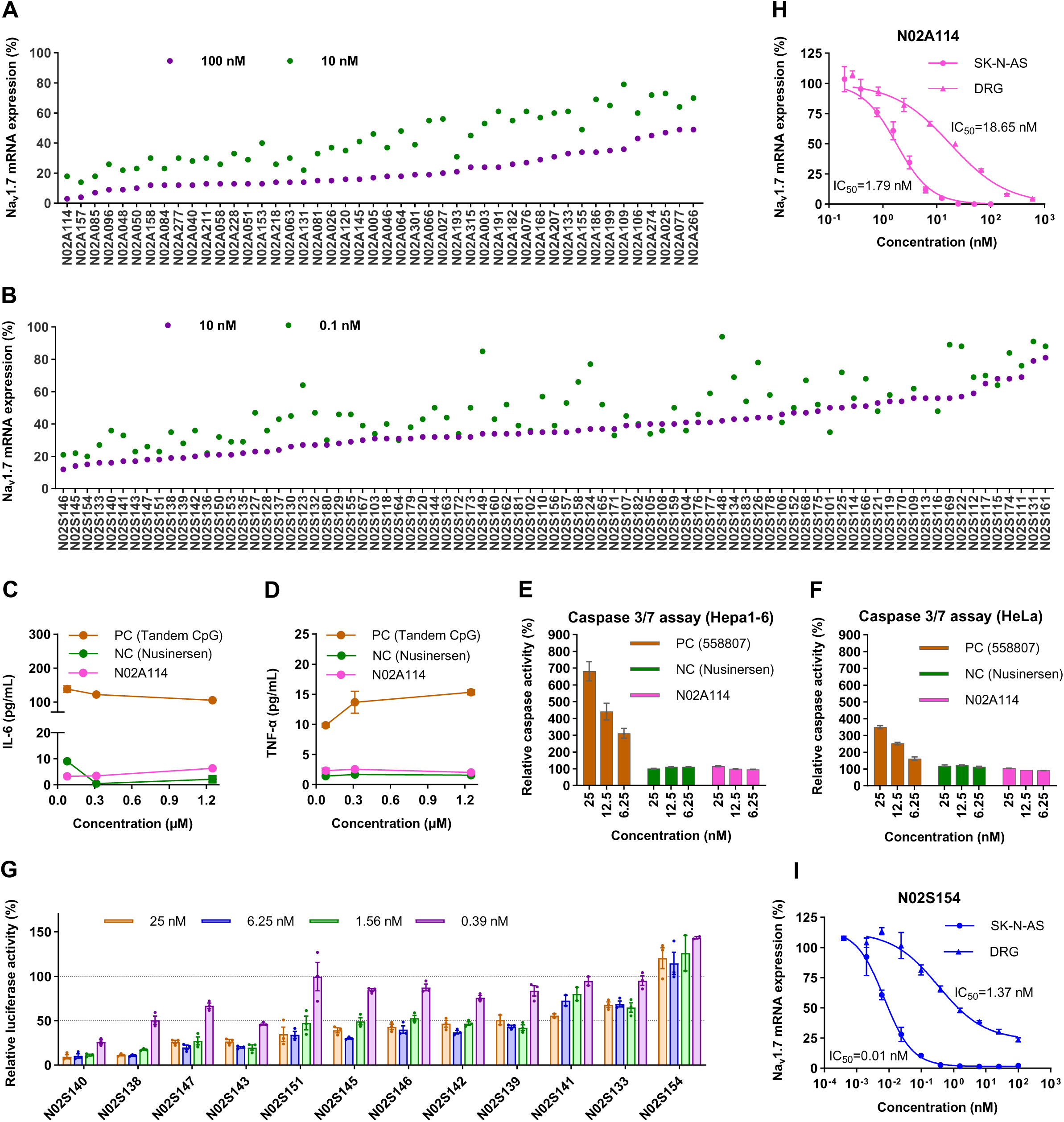
Screening and characterization of Nav1.7-targeting ASOs and siRNAs. **(A, B)** Primary screening in SK-N-AS cells. Nav1.7 mRNA levels were quantified by RT-qPCR 48 h after transfection with RNAiMAX. ASOs were tested at 10 and 100 nM and ranked by knockdown efficiency at 100 nM (**A)**. siRNAs were tested at 0.1 and 10 nM and ranked by knockdown efficiency at 10 nM (**B**). (**C, D)** Immunostimulatory potential of N02A114 in human PBMCs treated with 0.078, 0.31, or 1.25 μM ASO. IL-6 (**C**) and TNF-α (**D**) levels were measured after 24 h by ELISA. Nusinersen and tandem CpG oligonucleotides served as negative (NC) and positive (PC) controls, respectively. (**E, F)** Cytotoxicity assessment by caspase-3/7 activity in Hepa1-6 (**E**) and HeLa (**F**) cells 24 h after transfection with N02A114 (6.25, 12.5, or 25 nM). Nusinersen served as NC and the hepatotoxic ASO 558807 as PC. (**G)** Seed-mediated off-target activity of 12 siRNAs measured in HeLa cells using a dual-luciferase reporter assay at 0.39-25 nM. (**H, I)** Dose-response curves for N02A114 (**H**) and N02S154 (**I**) in SK-N-AS and rat DRG cells determined by RT-qPCR. n = 3. Data represent one of three independent experiments.

Next, because some gapmer ASOs can activate pattern recognition receptors (PRRs) and trigger innate immune responses^56^, we evaluated the potential immunostimulatory effects of the selected ASOs. To exclude candidates that might induce a human-specific proinflammatory response, such as increased secretion of interleukin-6 (IL-6) and tumor necrosis factor-α (TNF-α)^57^, freshly isolated human peripheral blood mononuclear cells (hPBMCs) were treated with each ASO at 0.31 μM and 1.25 μM, and cytokine levels were quantified by ELISA. A tandem CpG oligonucleotide served as the positive control (PC)^58^, while nusinersen was used as the negative control (NC). Only N02A063 induced significant IL-6 elevation (70–84 pg/mL vs 21–38 pg/mL for negative control), whereas the remaining 17 ASOs, including N02A114, showed minimal cytokine induction (3–6 pg/mL for IL-6; 1.58–2.56 pg/mL for TNF-α) comparable to negative control, indicating low immunogenicity (**Figure 1C, D; Figure S1N**).

Furthermore, in vitro cytotoxicity was assessed using a caspase 3/7 activity assay, a method strongly correlated with in vivo hepatic apoptosis and predictive of potential hepatotoxicity^59,60^. The 17 ASOs with minimal proinflammatory risk were transfected into murine Hepa1-6 and human HeLa cells at 25 nM, and caspase 3/7 activity was measured 24 h post-transfection. ASO 558807, previously validated as hepatotoxic in rats, served as the positive control (PC)^60^, while nusinersen was used as the negative control (NC). Baseline activity for negative control was ∼121% (Hepa1-6) and ∼117% (HeLa), while the positive control (ASO 558807) induced ∼215–248%. The relative caspase 3/7 activities for 17 ASOs ranged 82–138%, with N02A114 showing 92–116%, indicating negligible cytotoxicity (**Figure 1E, F; Figure S1O, P**). Collectively, N02A114 was selected as the optimal ASO candidate for subsequent in vivo studies.

Thirdly, siRNAs can induce off-target effects by partially binding non-target mRNAs through the seed region (nucleotides 2-8 at the 5’ end of the guide strand)^61^. Seed-dependent off-target effects were assessed using the psiCHECK-2 dual-luciferase assay^62^. Eleven of 12 candidates displayed dose-dependent off-target activity, whereas N02S154 maintained near-100% luciferase activity across 0.39–25 nM, indicating minimal off-target risk (**Figure 1G**).

Finally, dose-response validation in SK-N-AS and rat DRG cells showed N02A114 IC₅₀ of 1.79 nM and 18.65 nM, respectively, and N02S154 IC₅₀ of 0.01 nM and 1.37 nM, respectively (**Figure 1H, I**). These results demonstrate that N02A114 and N02S154 provide potent, selective Nav1.7 knockdown with negligible immunogenicity, hepatotoxicity, or off-target effects.

### In vivo efficacy of ASO N02A114 and siRNA N02S154 compared with pregabalin

To evaluate the in vivo inhibitory activity and analgesic effects of the lead ASO (N02A114) and siRNA (N02S154), dose-range finding (DRF) studies were first performed. Sprague–Dawley rats received a single intrathecal (IT) injection of ascending doses of N02A114 (0.1, 1.0, 3.0 mg/rat) or vehicle (aCSF). Fourteen days post-administration, RT-qPCR analysis revealed dose-dependent Nav1.7 knockdown in the spinal cord (s.c.) and dorsal root ganglia (DRGs): cervical s.c., 4–83%; thoracic s.c., 52–94%; lumbar s.c., 76–92%; DRGs, 9–63% (**Figure 2A-D**). Knockdown was strongest in the lumbar s.c., likely reflecting proximity to the IT injection site, while cervical, thoracic, and DRG regions showed moderate reductions.

**Figure 2.**
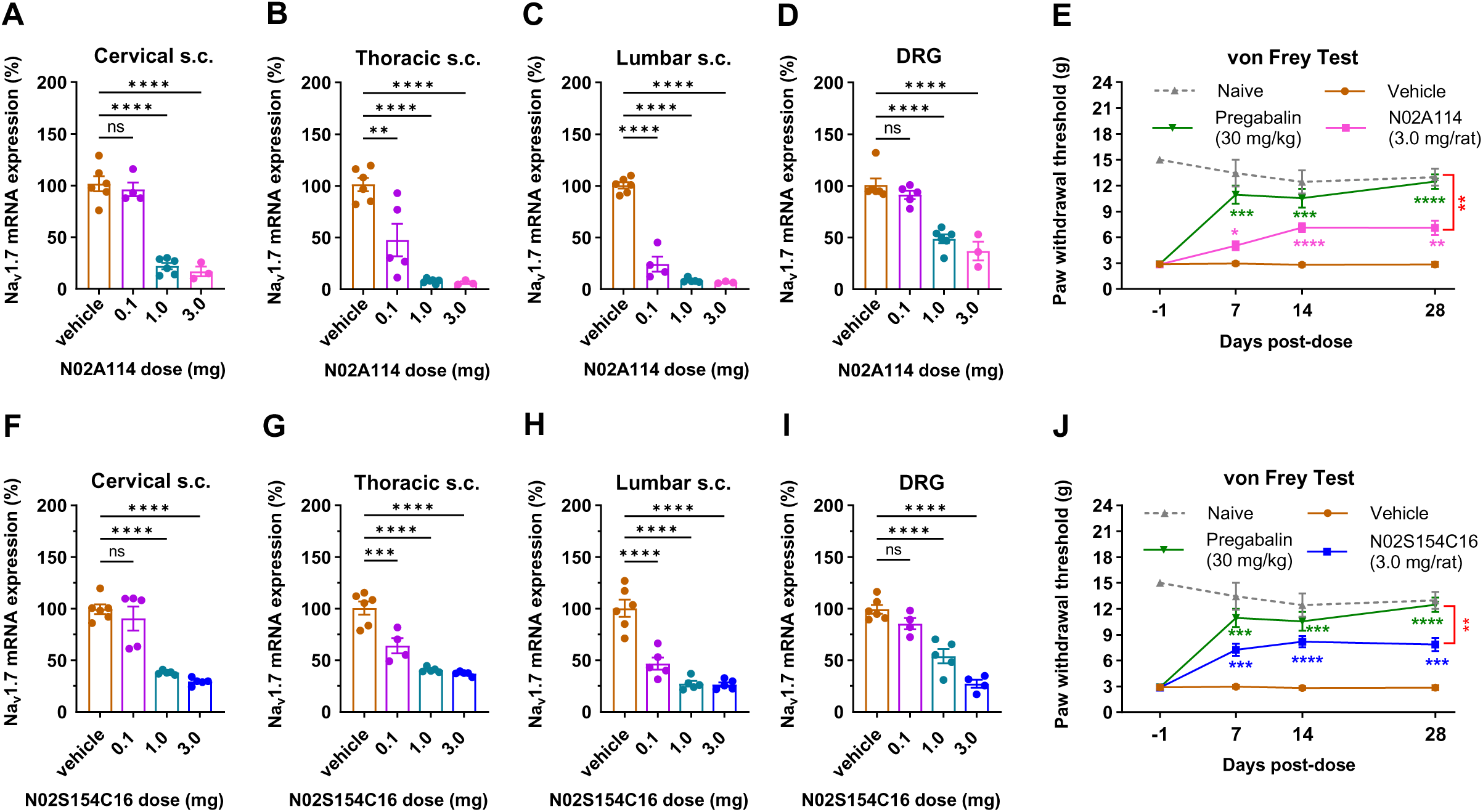
*In vivo* efficacy of N02A114 and N02S154. **(A-D)** Dose-range finding (DRF) study of N02A114. Rats received a single intrathecal (IT) injection of N02A114 (0.1, 1.0, or 3.0 mg/rat). Cervical (**A**), thoracic (**B**), and lumbar (**C**) spinal cord, and DRG (**D**) tissues were collected 14 days post-dose for Nav1.7 mRNA quantification by RT-qPCR. n = 6 animals for vehicle; n = 5 for 0.1 and 1 mg/rat; n = 3 for 3 mg/rat. Statistical significance was determined by one-way ANOVA with Dunnett’s multiple-comparisons test versus vehicle. (**E)** Mechanical sensitivity was assessed by von Frey testing on days 7, 14, and 28 in SNL rats following a single IT dose of N02A114 (3 mg/rat; day 0). Pregabalin (30 mg/kg) was administered by oral gavage 2 h before testing. The comparison between N02A114 and pregabalin at day 28 is indicated by the bracket. n = 8 animals per group for vehicle and pregabalin; n = 10 for N02A114; n = 3 for naive controls. Statistical analysis was performed using two-way ANOVA with Tukey’s multiple-comparisons test. **(F-I)** DRF study of N02S154C16. Rats received a single IT injection of N02S154C16 (0.1, 1.0, or 3.0 mg/rat). Cervical (**F**), thoracic (**G**), and lumbar (**H**) spinal cord, and DRG (**I**) tissues were collected 14 days post-dose for Nav1.7 mRNA quantification by RT-qPCR. n = 6 animals for vehicle and n = 5 per treatment group. Statistical significance was determined by one-way ANOVA with Dunnett’s multiple-comparisons test versus vehicle. (**J)** Mechanical allodynia was assessed by von Frey testing on days 7, 14, and 28 in SNL rats following a single IT dose of N02S154C16 (3 mg/rat; day 0). Pregabalin (30 mg/kg) was administered by oral gavage 2 h before testing. The comparison between N02S154C16 and pregabalin at day 28 is indicated by the bracket. n = 8 animals per group for vehicle and pregabalin; n = 13 for N02S154C16; n = 3 for naive controls. Statistical analysis was performed using two-way ANOVA with Tukey’s multiple-comparisons test. Data are presented as mean ± SEM. ns *P* > 0.05, \**P* < 0.05, \*\**P* < 0.01, \*\*\**P* < 0.001, \*\*\*\**P* < 0.0001.

Analgesic effects were assessed anti-nociceptive activity using the von Frey test in rats subjected to spinal nerve ligation (SNL), a well-established model of chronic pain characterized by tactile allodynia^63^. Rats in the treatment group received a single intrathecal (IT) injection of N02A114 at 3.0 mg/rat, and mechanical pain responses were measured on days 7, 14, and 28 post-dosing. A positive control group was orally administered pregabalin at 30 mg/kg two hours prior to each testing timepoint. Nav1.7 mRNA levels in the treatment group showed sustained knockdown, with reductions ranging from 96% to 87% in the lumbar spinal cord and from 87% to 53% in the DRGs (**Figure S2A, B**). In the von Frey assay, N02A114-treated rats exhibited significantly elevated paw withdrawal thresholds (PWTs) compared with vehicle controls at all timepoints (**Figure 2E**). The vehicle group had an average baseline PWT of 2.95 ± 0.23 g, whereas N02A114 increased the PWT to 5.31 ± 0.54 g (\**P* = 0.0137) on day 7. The analgesic effect further increased, reaching a peak PWT of 7.09 ± 0.47 g on day 14 (\*\*\*\**P* < 0.0001). By day 28, the PWT plateaued at 7.08 ± 0.86 g, significantly higher than the vehicle group (2.85 ± 0.28 g, \*\**P* = 0.0033), but still lower than the pregabalin group (12.49 ± 0.84 g, \*\**P* = 0.0019).

For N02S154, conjugation with a C16 moiety (N02S154C16) enhanced CNS delivery^64^. SD rats received a single IT injection at 0.1, 1.0, or 3.0 mg/rat. Fourteen days post-dose, Nav1.7 knockdown was dose-dependent: cervical s.c., 9–71%; thoracic s.c., 36–63%; lumbar s.c., 53–73%; DRGs, 15–73% (**Figure 2F-I**). The analgesic efficacy of N02S154C16 was further assessed in SNL rats, a single 3.0 mg/rat dose significantly alleviated tactile allodynia, with PWTs of 7.25 ± 0.69 g (day 7, \*\*\**P* = 0.0002), 8.18 ± 0.64 g (day 14, \*\*\*\**P* < 0.0001), and 7.87 ± 0.77 g (day 28, \*\*\**P* = 0.0001) (**Figure 2J**), corresponding to sustained Nav1.7 knockdown (lumbar s.c. 71–65%, DRGs 69–53%; **Figure S2C, D**). These values were significantly higher than those of the vehicle group but remained below the PWTs observed in the pregabalin group.

In summary, both N02A114 and N02S154 achieved substantial Nav1.7 suppression. However, neither matched the analgesic efficacy of pregabalin at any tested time point. These findings indicate that, despite the promise of ASO and siRNA platforms, single-molecule Nav1.7-targeting constructs are insufficient for optimal pain relief, highlighting the need for a more potent therapeutic strategy.

### Optimization of the ASO–siRNA Conjugation Strategy

To maximize Nav1.7 inhibition, we hypothesized that combining ASO and siRNA targeting the same transcript could synergistically enhance knockdown. We designed a series of ASO–siRNA conjugates (ASCs) covalently linked via a PEG6 linker, exploring three conjugation strategies: N02C0701, N02C0702, and N02C0703, differing in ASO-to-siRNA stoichiometry and attachment site (**Figure 3A**). SK-N-AS cells were transfected with 0.1, 1, and 10 nM of each ASC, and Nav1.7 mRNA levels were measured by RT-qPCR 48 h post-transfection. As expected, single-molecule siRNA N02S154 exhibited higher knockdown than N02A114 and served as reference. At 10 nM, N02C0701 (99.90%, \*\*\**P* = 0.0002) and N02C0702 (99.94%, \*\*\**P* = 0.0001) achieved significantly greater knockdown than N02S154 (97.39%), whereas N02C0703 (97.01%) showed no improvement. At 1 nM, only N02C0702 (97.16%) significantly outperformed N02S154 (88.09%, \**P* = 0.0205). At 0.1 nM, knockdown by all ASCs was comparable or slightly lower than N02S154, demonstrating dose-dependent effects (**Figure 3B**).

**Figure 3.**
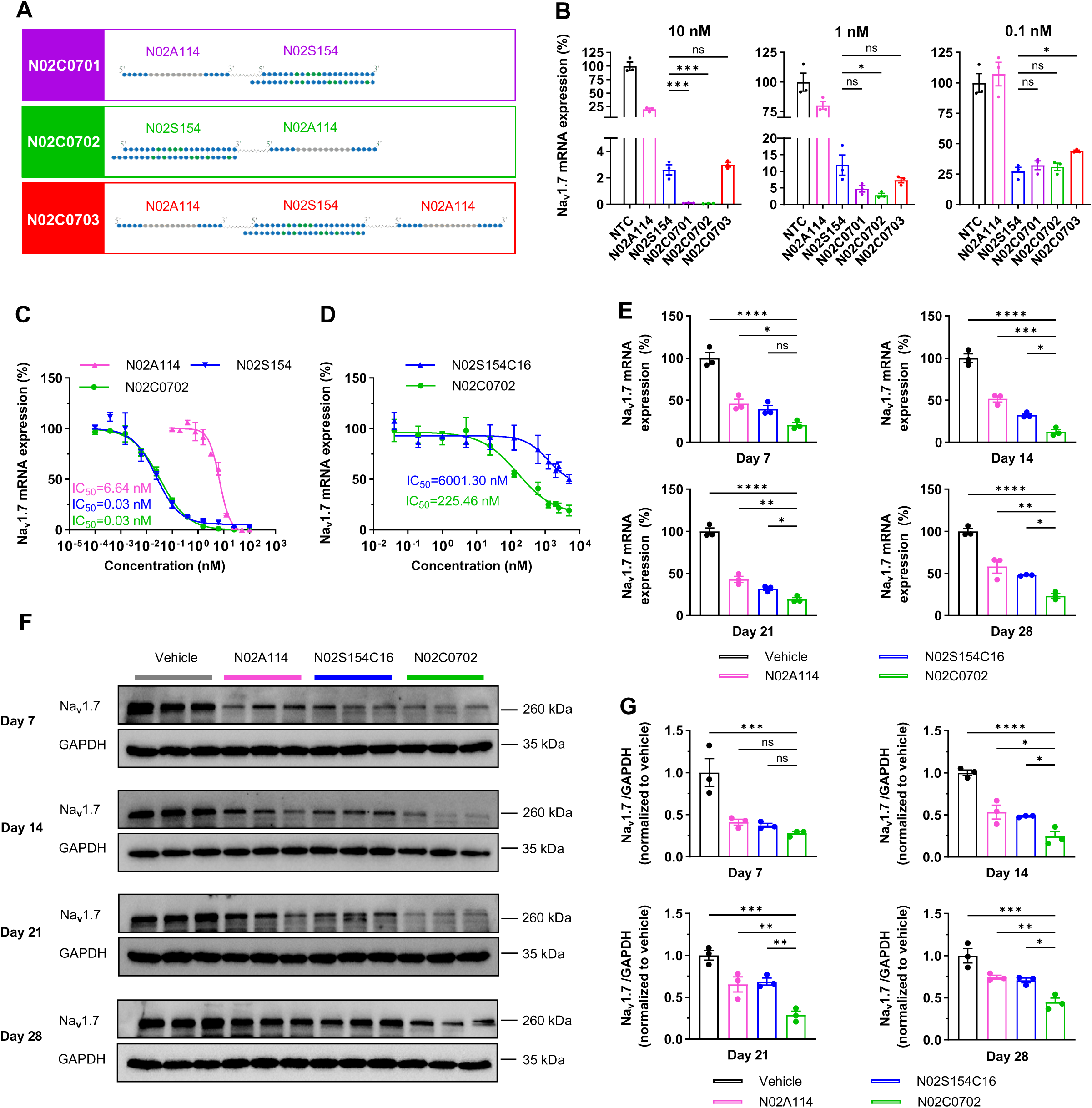
Characterization of ASO–siRNA conjugates and comparison with corresponding ASO or siRNA counterparts. **(A)** Schematic representation of the conjugate designs. N02C0701 (purple): the 3’ end of the ASO is linked to the 5’ terminus of the siRNA sense strand (SS) via a PEG6 linker. N02C0702 (green): the 5’ end of the ASO is linked to the 3’ terminus of the siRNA SS via a PEG6 linker. N02C0703 (red): the siRNA SS is conjugated to two identical ASOs, with the 5’ terminus of the siRNA linked to the 3’ end of one ASO and the 3’ terminus linked to the 5’ end of the second ASO. **(B)** SK-N-AS cells were transfected with N02A114 (ASO), N02S154 (siRNA), or the corresponding conjugates at 0.1, 1, or 10 nM for 48 h. Relative Nav1.7 mRNA levels were measured by RT-qPCR. Statistical significance was determined by one-way ANOVA with Dunnett’s multiple-comparisons test using N02S154 as the reference. n = 3 independent experiments. **(C)** Dose–response curves in SK-N-AS cells comparing knockdown activity of the ASO-siRNA conjugate N02C0702 with the corresponding ASO (N02A114) and siRNA (N02S154C16) following transfection. n = 3 independent experiments. **(D)** Dose–response curves in rat DRG neurons following free uptake, comparing knockdown activity of N02C0702 with N02S154C16. n = 3 experimental replicates. (**E, F)** Effects of intrathecal administration of N02A114 (1.5 mg/rat), N02S154C16 (3.0 mg/rat), or N02C0702 (4.5 mg/rat) on Nav1.7 mRNA (**E**) and protein (**F**) levels in DRG tissues at days 7, 14, 21, and 28 post-dose. n = 3 animals per group. **(G)** Quantification of Nav1.7 protein levels in rat DRG tissues. Statistical significance for **E** and **G** was determined by one-way ANOVA with Dunnett’s multiple-comparisons test using N02C0702 as the reference group. Data are presented as mean ± SEM. ns *P* > 0.05, \**P* < 0.05, \*\**P* < 0.01, \*\*\**P* < 0.001, \*\*\*\**P* < 0.0001.

To validate the optimal conjugation pattern, additional ASCs were synthesized using alternative ASOs (N02A157, N02A114) and siRNAs (N02S154, N02S133): N02C0501–0503, N02C1201–1203, and N02C1401–1403 (**Figure S3A-C; Table S3**). At 10 nM, all ASCs achieved significantly higher knockdown than siRNA alone: N02C0501 (99.17%, \*\**P* = 0.0084), N02C0502 (99.76%, \*\**P* = 0.0014), N02C0503 (99.10%, \**P* = 0.0105) versus N02S154; N02C1201 (99.13%), N02C1202 (99.42%), N02C1203 (96.66%, \*\*\*\**P* < 0.0001); N02C1401 (99.73%) and N02C1402 (99.73%, \*\*\**P* = 0.0003) versus N02S133 (93.06%) (**Figure S3D-F**). These results demonstrate that covalent ASO-siRNA conjugation significantly enhances Nav1.7 knockdown relative to single-molecule siRNA, with N02C0702 emerging as the most potent conjugation strategy for further in vitro and in vivo evaluation.

Across all ASCs tested, equimolar conjugates consistently outperformed their 2:1 counterpart. Specifically, N02C0701/N02C0702 exceeded N02C0703 at 10 nM and 0.1 nM (**Figure 3B**); N02C0501/N02C0502 outperformed N02C0503 at 1 nM and 0.1 nM (**Figure S3D**); N02C1201/N02C1202 surpassed N02C1203 at 10 nM (**Figure S3E**); and N02C1401/N02C1402 exceeded N02C1403 across 10 nM, 1 nM, and 0.1 nM (**Figure S3F**). Conjugation to the 3’-end of the siRNA sense strand was generally more favorable than 5’-end attachment, yielding consistently higher or equivalent knockdown. Among all designs, N02C0702, representing ASO tethered to the 3’-end in an equimolar configuration, achieved the highest Nav1.7 knockdown and was selected for subsequent in vivo analgesic evaluation.

### Enhanced Nav1.7 knockdown by N02C0702 in vitro and in vivo

Next, we compared the in vitro knockdown potency of N02C0702 with its ASO and siRNA counterparts by transfecting SK-N-AS cells with ascending doses (0.0001-100 nM) and calculating IC₅₀ values for Nav1.7 mRNA reduction. N02C0702 (IC₅₀ = 0.03 nM) exhibited ∼200-fold greater potency than N02A114 and was comparable to N02S154 (**Figure 3C**). Notably, the maximal knockdown of N02C0702 (0.14% mRNA remaining) exceeded that of N02S154 (3.01% remaining) and approached N02A114 (0.38% remaining), indicating synergistic enhancement when ASO and siRNA target the same transcript.

To confirm the ASO-mediated self-delivery capacity of ASCs, DRG cells were treated via free uptake with N02C0702 or N02S154C16 at doses ranging from 0.04 nM to 5 μM. 72 hours later, Nav1.7 mRNA knockdown was assessed. N02C0702 entered cells efficiently and achieved at least 25-fold greater potency than N02S154C16 (IC₅₀ = 225.46 nM vs. 6001.30 nM) (**Figure 3D**).

The in vivo potency of N02C0702 was then evaluated relative to its ASO and siRNA counterparts. Rats received a single IT injection of N02C0702 (4.5 mg/rat), N02A114 (1.5 mg/rat), or N02S154C16 (3.0 mg/rat) at nearly equimolar doses. Nav1.7 mRNA and protein levels were quantified by RT-qPCR and western blot at days 7, 14, 21, and 28 post-dosing. In DRG tissues, qPCR analysis confirmed that N02C0702 induced significantly greater Nav1.7 mRNA knockdown compared with N02A114 or N02S154 at most tested time points (\**P* < 0.05) (**Figure 3E**). Specifically, N02A114 reduced Nav1.7 mRNA by 54%, 48%, 57%, and 42% on days 7, 14, 21, and 28, respectively, while N02S154 achieved 61%, 68%, 68%, and 52%. In contrast, N02C0702 produced reductions of 80%, 88%, 81%, and 77% over the same period. Except for N02A114 at days 14 and 28, all treatments maintained >50% mRNA suppression, a threshold reported as essential for robust analgesia via Nav1.7 inhibition. Consistent with mRNA results, western blot analysis demonstrated that N02C0702 suppressed Nav1.7 protein expression more effectively than N02A114 or N02S154 from day 14 through day 28 (**Figure 3F, G**; \**P* < 0.05). Quantification showed N02C0702 maintained >70% protein reduction from days 7 to 21, decreasing slightly to 55% at day 28. In comparison, N02S154 inhibition declined from 63% to 30%, and N02A114 from 59% to 26% over the same period. In the spinal cord, Nav1.7 mRNA and protein levels were largely similar across the N02C0702, N02S154, and N02A114 groups at most time points (**Figure S4A-C**).

Overall, all three oligonucleotide treatments significantly reduced Nav1.7 expression in DRG and spinal cord tissues relative to vehicle controls. Notably, N02C0702 achieved the most robust and sustained knockdown, with consistent concordance between mRNA and protein levels in DRGs.

### N02C0702 shows superior and prolonged analgesic effects

To evaluate the therapeutic efficacy and duration of action of N02C0702, a dose-range finding (DRF) study was performed. Rats received a single IT injection of N02C0702 at 0.15, 1.5, or 4.5 mg/rat, and Nav1.7 mRNA levels in DRGs and spinal cord were measured by RT-qPCR on day 14. Dose-dependent knockdown was observed across all regions: cervical s.c. (39%, 78%, 79%), thoracic s.c. (53%, 71%, 74%), lumbar s.c. (69%, 81%, 85%), and DRGs (34%, 61%, 77%) (**Figure 4A**), demonstrating robust suppression in DRGs, the primary tissues associated with nociception.

**Figure 4.**
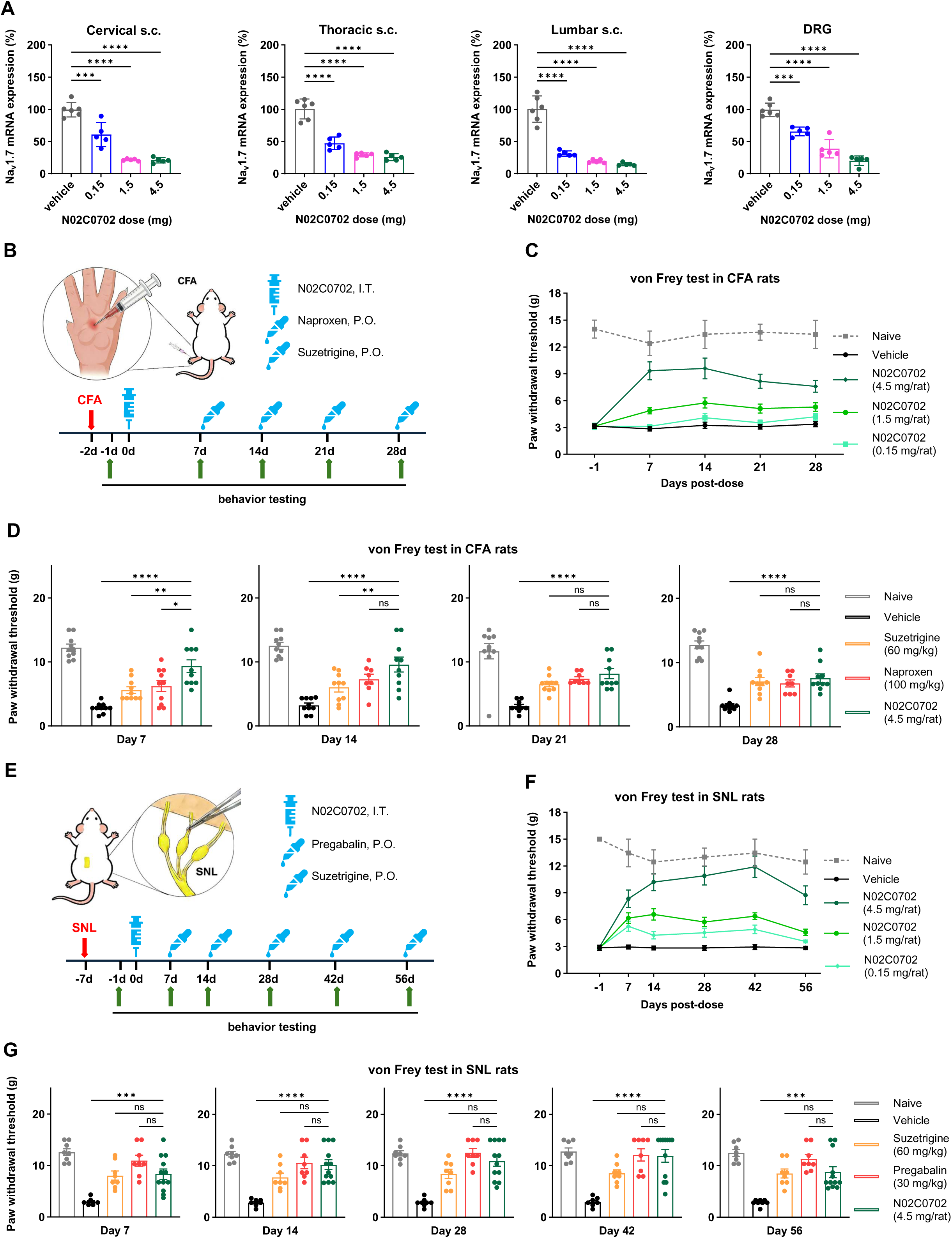
Analgesic effects of N02C0702 in CFA and SNL rat models. **(A)** Dose-range finding (DRF) study of N02C0702. Rats received a single intrathecal (IT) injection of N02C0702 (0.15, 1.5, or 4.5 mg/rat). Cervical, thoracic, and lumbar spinal cord and DRG tissues were collected 14 days post-dose for Nav1.7 mRNA quantification by RT-qPCR. n = 5 animals per treatment group and n = 6 for vehicle. Statistical significance was determined by one-way ANOVA with Dunnett’s multiple-comparisons test versus vehicle. **(B)** Experimental design for evaluating the effects of intrathecal N02C0702 and orally administered suzetrigine or naproxen on pain-related behaviors in the CFA-induced rat model of inflammatory pain. **(C)** Dose-dependent effects of N02C0702 (0.15, 1.5, or 4.5 mg /rat) on paw withdrawal thresholds (PWTs), measured by von Frey testing on days 7, 14, 21, and 28 after CFA induction. **(D)** Comparison of PWTs among N02C0702 (4.5 mg/rat), suzetrigine (60 mg/kg), naproxen (100 mg/kg), vehicle, and naive groups at indicated time points. For c, d, n = 10 animals per group. Statistical significance was determined by one-way ANOVA with Dunnett’s multiple-comparisons test using N02C0702 as the reference group. **(E)** Experimental design for evaluating the effects of intrathecal N02C0702 and orally administered suzetrigine or pregabalin in the spinal nerve ligation (SNL) rat model of neuropathic pain. **(F)** Dose-dependent effects of N02C0702 (0.15, 1.5, or 4.5 mg/rat) on PWTs measured by von Frey testing on days 7, 14, 28, 42, and 56 after SNL surgery. **(G)** Comparison of PWTs among N02C0702 (4.5 mg/rat), suzetrigine (60 mg/kg), pregabalin (30 mg/kg), vehicle, and naïve groups at indicated time points. For f, g, naïve, vehicle, suzetrigine, and pregabalin groups: n = 8 animals per group; N02C0702 group: n = 12 animals. Statistical significance was determined by one-way ANOVA with Dunnett’s multiple-comparisons test using N02C0702 as the reference group. Data are presented as mean ± SEM. ns P > 0.05, \**P* < 0.05, \*\**P* < 0.01, \*\*\**P* < 0.001, \*\*\*\**P* < 0.0001.

We next assessed anti-nociceptive efficacy in the CFA model of inflammatory pain. N02C0702 administered at the same doses produced dose-dependent analgesia, as measured by von Frey testing at 1-, 2-, 3-, and 4-weeks post-dose. High-dose N02C0702 (4.5 mg/rat) significantly elevated paw withdrawal thresholds (PWTs) relative to vehicle-treated rats, achieving 9.34 ± 0.10 g at day 7 and 9.59 ± 1.16 g at day 14 (**Figure 4B-D**). These values were superior to naproxen (6.24 ± 0.87 g) and suzetrigine (5.61 ± 0.52 g) at day 7 (*P < 0.05). The analgesic effect of N02C0702 persisted throughout the 4-week evaluation period, reaching comparable PWTs to positive controls at later time points.

In the SNL model of neuropathic pain, rats received a single IT injection of N02C0702 (0.15, 1.5, or 4.5 mg/rat), with pregabalin (30 mg/kg) and suzetrigine (60 mg/kg) as oral positive controls. Mechanical allodynia was monitored via von Frey test on days 7, 14, 28, 42, and 56 post-dose (**Figure 4E-G**). All N02C0702-treated groups displayed dose- and time-dependent increases in PWTs relative to vehicle (2.83–2.96 g). In the high-dose group, PWTs rose from 8.33 ± 0.99 g on day 7 to a peak of 11.90 ± 1.21 g on day 42, then declined slightly to 8.74 ± 1.06 g on day 56. Medium- and low-dose groups exhibited similar trends with lower magnitude. Across the study period, high-dose N02C0702 achieved analgesic efficacy comparable to pregabalin (10.54–12.49 g) and suzetrigine (7.73–8.52 g), with no significant differences at most time points (*P* > 0.05) (**Figure 4G**).

Thermal hyperalgesia assessed by the Hargreaves test mirrored mechanical outcomes. High-and medium-dose N02C0702 increased paw withdrawal latencies (PWLs) from baseline 8.32 s to 16.14 ± 1.24 s and 13.03 ± 0.67 s at day 28, respectively, gradually declining by day 56. Low-dose rats showed minimal effect. PWLs in pregabalin- and suzetrigine-treated rats reached 15.70 ± 0.35 s and 15.85 ± 0.35 s, indicating that a single high-dose N02C0702 achieved analgesic potency comparable to both positive controls over 56 days (**Figure S5A, B**).

Collectively, N02C0702 exhibited a robust dose-response relationship and reproducible analgesic efficacy across both inflammatory and neuropathic pain models. A single IT administration produced rapid onset of analgesia from day 7, which persisted for at least 28 days in CFA-induced pain and 56 days in SNL-induced pain. These results highlight N02C0702 as a potent and durable ASC candidate with strong translational potential for the treatment of chronic pain.

### Off-target and genome-wide specificity of N02C0702

To evaluate potential off-target effects, genome-wide RNA sequencing was performed in SK-N-AS cells 24 h after transfection with N02C0702 at 0.1, 10, and 30 nM (**Figure 5A**). Genes with an adjusted p-value < 0.05 and (Log2FC) > 1 were considered significantly regulated. At 0.1 nM, SCN9A-the intended target-was the only outlier (Log2FC = −1.6, 67% knockdown, P< 10^-4), consistent with RT-qPCR (70% reduction), confirming on-target specificity. At 10 nM (about 300-fold IC_50_), only SLC39A10 showed modest downregulation (Log2FC = −1.05). At a supratherapeutic 30 nM, 36 genes were differentially expressed (9 downregulated, 24 upregulated), with modest fold changes; gene ontology analysis revealed enrichment among upregulated genes for chromatin structure and chloride transporter activity, likely reflecting high-dose or synthesis-related effects rather than biologically relevant off-targets.

**Figure 5.**
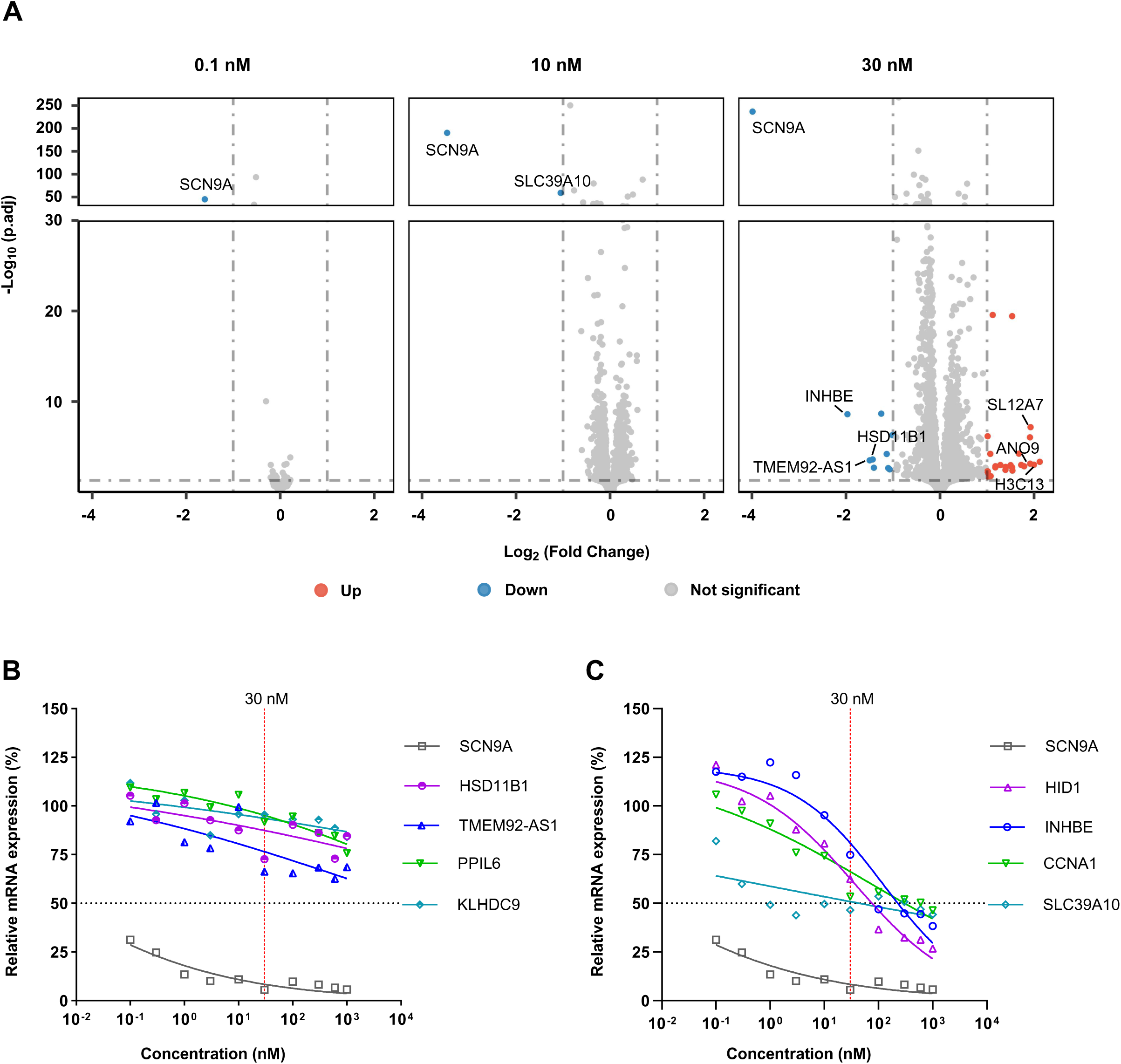
Off-target assessment of N02C0702 by RNA sequencing and validation of candidate genes by RT–qPCR. **(A)** Volcano plot of RNA-seq data from SK-N-AS cells transfected with N02C0702 (0.1, 10, or 30 nM) for 24 h. The x-axis represents the log₂ fold change between N02C0702-treated and vehicle-treated groups, and the y-axis represents the −log₁₀(P) value. Blue dots indicate downregulated genes (log₂ fold change < −1 and P < 0.05), red dots indicate upregulated genes (log₂ fold change > 1 and false discovery rate (FDR) < 0.05), and gray dots indicate non-significant genes (|log₂ fold change| ≤ 1 or P ≥ 0.05). n = 3 experimental replicates. (**B, C)** RT-qPCR validation of candidate off-target genes identified by RNA-seq. Dose-response curves were generated following transfection with N02C0702 at concentrations ranging from 0.1 to 1,000 nM. The 1,000× IC₅₀ value of N02C0702 is indicated by a red dotted line. n = 3 experimental replicates.

Genes exhibiting maximal response and Hill curve fitting were further examined by RT-qPCR. Differential expressed gene SLC39A10 in 10 nM group, along with 7 genes (TMEM92-AS1, HSD11B1, PPIL6, KLHDC9, INHBE, HID1, CCNA1) in 30 nM group (excluding 2 without official gene symbols), were further validated by RT-qPCR (primers in **Table S4**). Among eight candidates, four (TMEM92-AS1, HSD11B1, PPIL6, KLHDC9) showed no substantial reduction (>50%) even at 1000 nM (>30,000-fold SCN9A IC_50_) (**Figure 5B**). Three genes (CCNA1, HID1, INHBE) displayed dose-dependent downregulation with IC_50_ values of 304.88, 57.97, and 228.35 nM, >1,000-fold higher than SCN9A, indicating minimal clinical risk (**Figure 5C**). SLC39A10 reached a plateau at ∼50% knockdown even at 1000 nM, consistent with sequencing data (Log2FC = −0.96). SLC39A10 encodes a zinc transporter with no established disease association. Human population data from gnomAD v2.1.1 identified four heterozygous loss-of-function variants without uniform disease phenotypes, suggesting that partial suppression (∼50%) is likely non-pathogenic.

Overall, N02C0702 demonstrated robust on-target efficacy, long-lasting activity, and minimal off-target effects even at concentrations 1,000-fold above the SCN9A IC_50_, supporting its favorable safety profile and potential as a therapeutic candidate for chronic pain, with a pattern applicable to other indications.

## DISCUSSION

Durable, target-specific, non-addictive analgesics remain a critical unmet need. We report an oligonucleotide scaffold (ASC) that covalently links an ASO and an siRNA via a PEG6 linker, enabling dual targeting of Nav1.7. The lead ASC, N02C0702, achieved robust knockdown of Nav1.7 mRNA and protein in DRG and spinal cord, producing potent, long-lasting analgesia in vivo. These findings highlight ASCs as a promising platform for non-addictive pain therapy and reinforce Nav1.7 as a key analgesic target.

Sequences for the ASO and siRNA components were designed using rigorous algorithms. Overlapping regions were prioritized to inhibit both Nav1.7 splice isoforms^65^, and sequences with ≤3 mismatches to other genes, including the eight other Nav channels, were excluded. Conserved regions across humans, rodents, and non-human primates were selected to enhance translational relevance, and sequences containing SNP loci were omitted to maximize population applicability. Rational sequence design minimized off-target effects, as verified by RNA-sequencing, and contributed to the favorable initial in vivo safety of ASCs. Cross-species efficacy was confirmed in human and rat cells prior to in vivo validation^66^. Standard chemical modifications (PS, 2’-OMe, 2’-MOE) further supported preclinical and clinical safety. Although monomeric ASOs or siRNAs alone were insufficient for robust analgesia, their optimized design ensured the safety of the final ASC formulation.

ASCs achieved superior Nav1.7 knockdown compared with monomeric oligonucleotides (**Figure 3E-G**), reflecting the synergistic action of complementary mechanisms: ASOs induce RNase H–mediated mRNA degradation, whereas siRNAs engage RISC-mediated RNA interference^67–71^. Structurally analogous designs, such as asymmetric siRNAs^72^ and SCAD scaffolds^73^, have demonstrated efficient delivery via PS-modified single-stranded structures. Consistent with these observations, the dual-targeting ASC design amplifies therapeutic efficacy while leveraging the intrinsic self-delivery properties of PS-modified ASOs.

Analgesic effects in CFA and SNL models closely mirrored reductions in Nav1.7 mRNA and protein, with efficacy evident as early as day 7, earlier than previously reported^74^. In the CFA model, analgesic efficacy remained stable over 28 days, whereas in SNL it progressively increased and persisted through day 42, reflecting the distinct pathophysiology of inflammatory versus neuropathic pain. In the CFA model, N02C0702 matched naproxen efficacy, yet avoided NSAID-associated toxicities; notably, naproxen caused mortality in 2 of 8 rats. In the SNL model, ASC N02C0702 produced analgesia comparable to pregabalin but with a single administration sustaining effects for over 56 days and a reduced risk of adverse events. Across both models, ASC N02C0702 also outperformed suzetrigine, a Nav1.8-targeting agent. Some discrepancies between mRNA and protein knockdown in spinal cord may reflect limited Western blot sensitivity or Nav1.7 trafficking in sensory neurons. In contrast, DRG knockdown was consistent, underscoring its central role in nociceptive signaling. Whether spinal Nav1.7 directly contributes to pain transmission remains to be clarified.

Currently approved CNS oligonucleotide therapies are ASOs—Tofersen for ALS^75^ and Nusinersen for SMA^76^—targeting rare diseases, while no siRNA drugs have yet reached the CNS clinic. In hepatocytes, siRNA therapeutics exhibit potent, durable efficacy, with GalNAc conjugation achieving multiple clinical approvals^77^. ASCs provide a complementary strategy to deliver duplex RNAs to CNS and extrahepatic tissues. By integrating ASO and siRNA components, ASCs combine distinct silencing mechanisms while retaining PS-mediated delivery efficiency. The PEG6 conjugation is compatible with solid-phase synthesis and scalable manufacturing, supporting broad therapeutic potential. Reprogramming sequences could adapt ASCs to diverse targets.

Single-target inhibition is effective in monogenic diseases but limited in polygenic or adaptive disorders due to compensatory pathways. Multi-target strategies are increasingly pursued: Arrowhead’s ARO-DIMER-PA silences PCSK9 and APOC3 for mixed dyslipidemia (Phase 1/2a, NCT07223658)^78^, and Corsera’s COR-1004 targets PCSK9 and AGT in atherosclerosis (NCT07229118)^79^. The ASC scaffold can be converted into bifunctional constructs by reprogramming either oligonucleotide, circumventing competition for shared enzymatic pathways seen in homotypic conjugates (e.g., siRNA–siRNA). In analgesic development, ASCs allow concurrent targeting of Nav1.7 and Nav1.8, modulating both initiation and propagation of pain for potentially enhanced efficacy.

Despite the promising efficacy of ASC N02C0702 and the absence of observable weight loss in treated rats, further studies are needed to confirm long-term safety and support clinical translation. Head-to-head comparisons with other conjugates, such as di-siRNA^80^ or SCAD^73^, remain pending. Nonetheless, these results demonstrate ASCs as a versatile platform capable of potent, durable gene silencing in vitro and in vivo, with potential applicability to additional targets for diverse therapeutic indications.

## ACKNOWLEDGMENTS

We thank all staff at Shanghai Nyuen Biotechnology Co., Ltd. and the Kunming Institute of Zoology, Chinese Academy of Sciences, for their contributions to routine research and management. This work was supported by the Science and Technology Commission of Shanghai Municipality (B.R.), the National Natural Science Foundation of China (NSFC, 32571205 to G.W.), and Department of Science and Technology of Yunnan Province supported the “Xing Dian Talent Program” (G.W.).

## AUTHOR CONTRIBUTIONS

B.R. and G.W. conceptualized and supervised the research and wrote the manuscript. B.R. and C.Y. designed the experiments and analyzed the results. Sequence design and bioinformatic analyses were performed by B.R. and H.C. Cellular biology and biochemistry experiments were conducted by Y.L., B.F., and L.G., who also analyzed the data. Animal experiments were led by Z.Y., with assistance from S.L., Z.C., Q.Y., P.Y., X.Q., and W.Y. All authors reviewed and approved the final manuscript and contributed insights throughout the development of this work.

## Supplemental Figures

**Figure S1.**
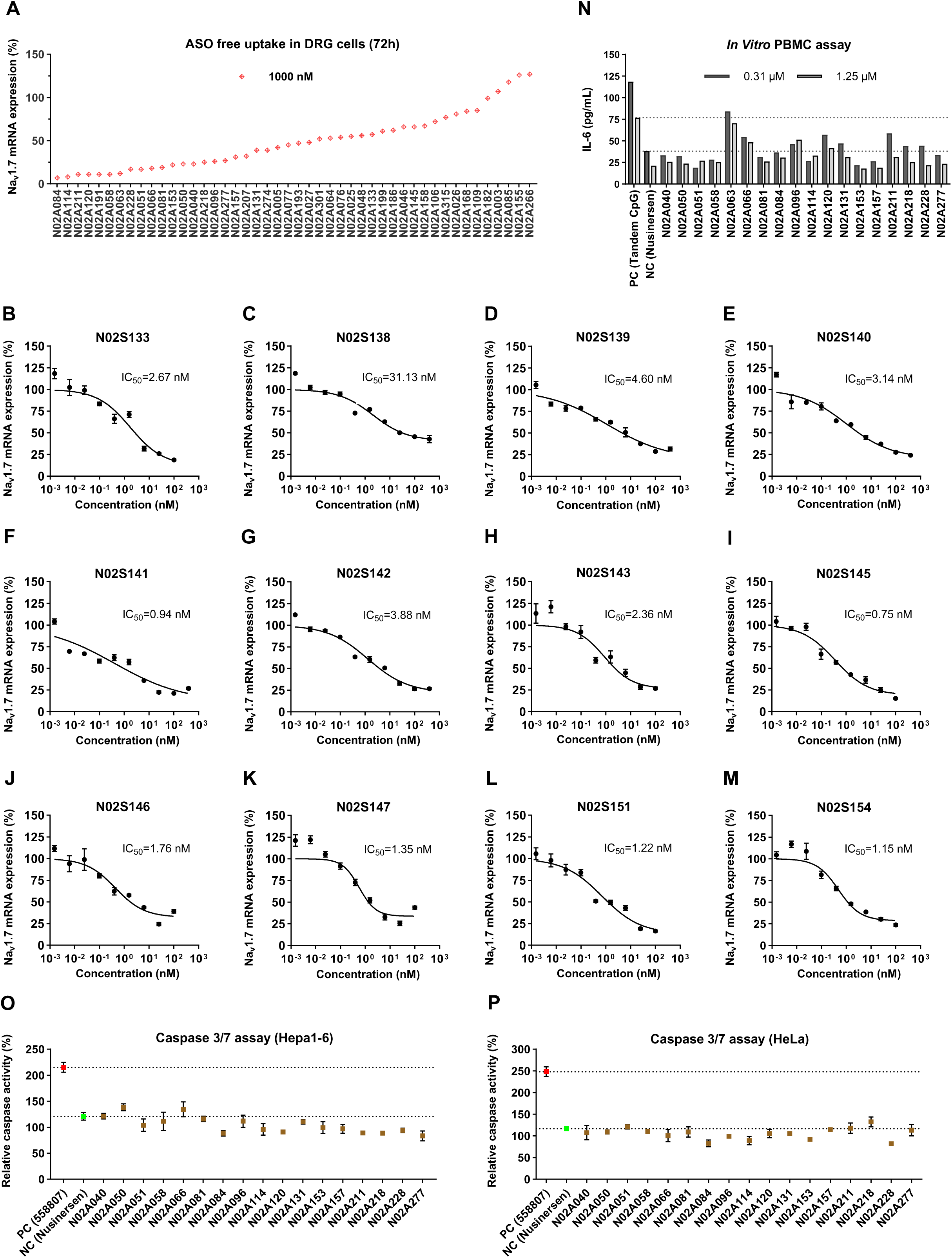
In vitro potency of ASOs and siRNAs in DRG cells and safety assessment of selected ASOs. **(A)** Screening of ASOs in rat DRG cells at a concentration of 1,000 nM. ASOs were delivered to DRG cells by free uptake, and Nav1.7 mRNA levels were measured 72 h after treatment by RT-qPCR. ASOs were ranked according to their knockdown activity. n = 3 experimental replicates. **(B-M)** Dose-response curves for 12 siRNAs in rat DRG cells. Cells were treated via free uptake and Nav1.7 mRNA levels were quantified by RT-qPCR. n = 3 experimental replicates. **(N)** Immunostimulatory assessment of selected ASOs in human peripheral blood mononuclear cells (PBMCs). Eighteen ASOs were incubated with PBMCs at 0.31 or 1.25 μM for 24 h, and IL-6 levels were measured using a sandwich ELISA. Nusinersen and tandem CpG oligodeoxynucleotides served as the negative control (NC) and positive control (PC), respectively. n = 3 experimental replicates. **(O, P)** Caspase 3/7 activity in Hepa1-6 (**O**) and Hela (**P**) cells was measured at 24 h after transfection of 17 selected ASOs at 25 nM. A reported sequence (558807) served as a PC and Nusinersen served as a NC. n = 3 experimental replicates. Data are presented as mean ± SEM.

**Figure S2.**
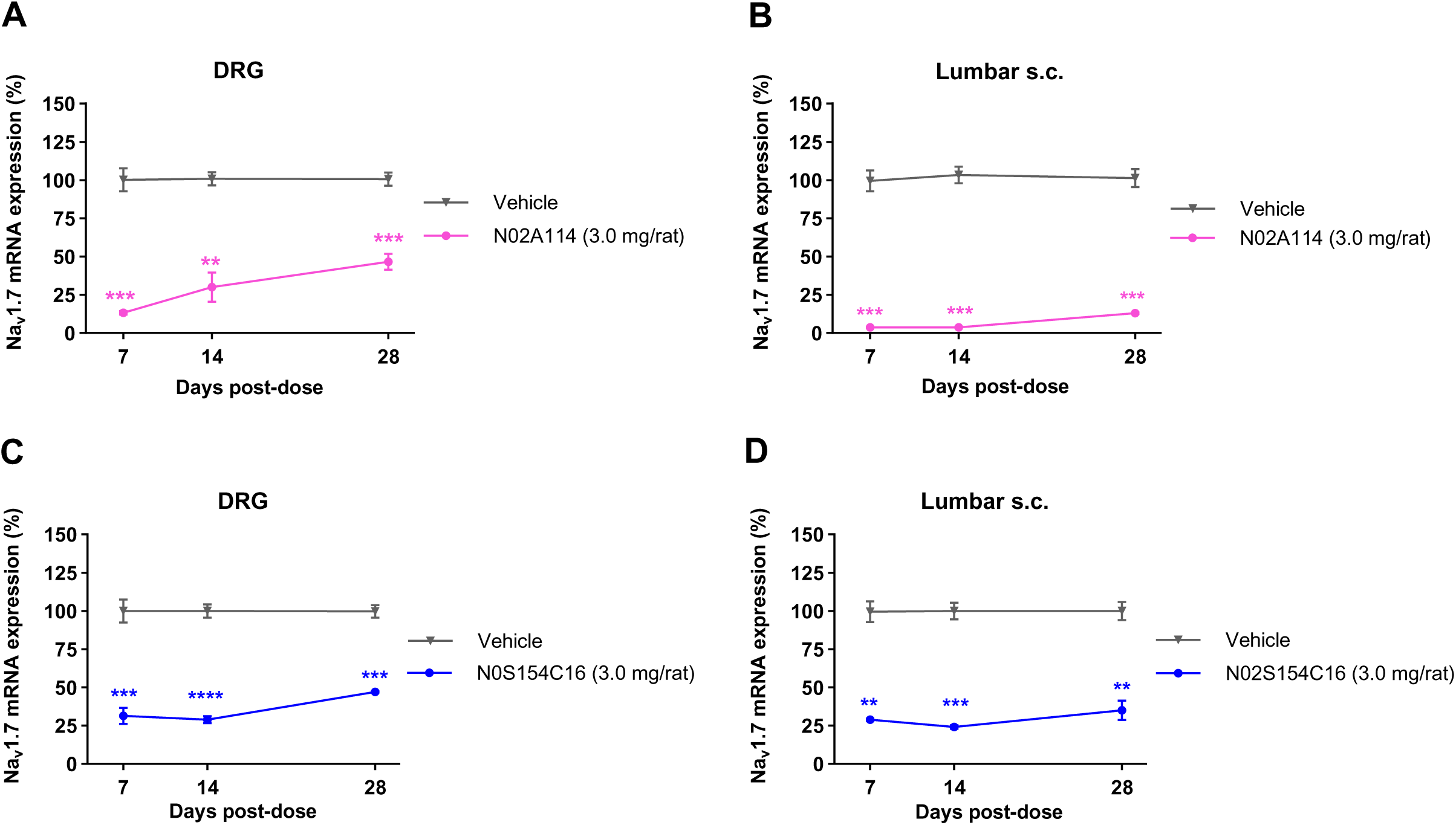
In vivo potency of N02A114 and N02S154C16. **(A, B)** Rats received a single intrathecal (IT) bolus injection of N02A114 (3.0 mg/rat). DRG (**A**) and lumbar spinal cord (**B**) tissues were collected at the indicated time points post-dose for Nav1.7 mRNA quantification by RT-qPCR. n = 5 animals per group. Statistical significance was determined by two-way ANOVA with Tukey’s multiple-comparisons test. **(C, D)** Rats received a single IT bolus injection of N02S154C16 (3.0 mg/rat). DRG (**C**) and lumbar spinal cord (**D**) tissues were collected at the indicated time points post-dose for Nav1.7 mRNA quantification by RT-qPCR. n = 5 animals for the vehicle group and n = 3 animals for the N02S154C16-treated group. Data are presented as mean ± SEM. ns *P* > 0.05, \**P* < 0.05, \*\**P* < 0.01, \*\*\**P* < 0.001, \*\*\*\**P* < 0.0001.

**Figure S3.**
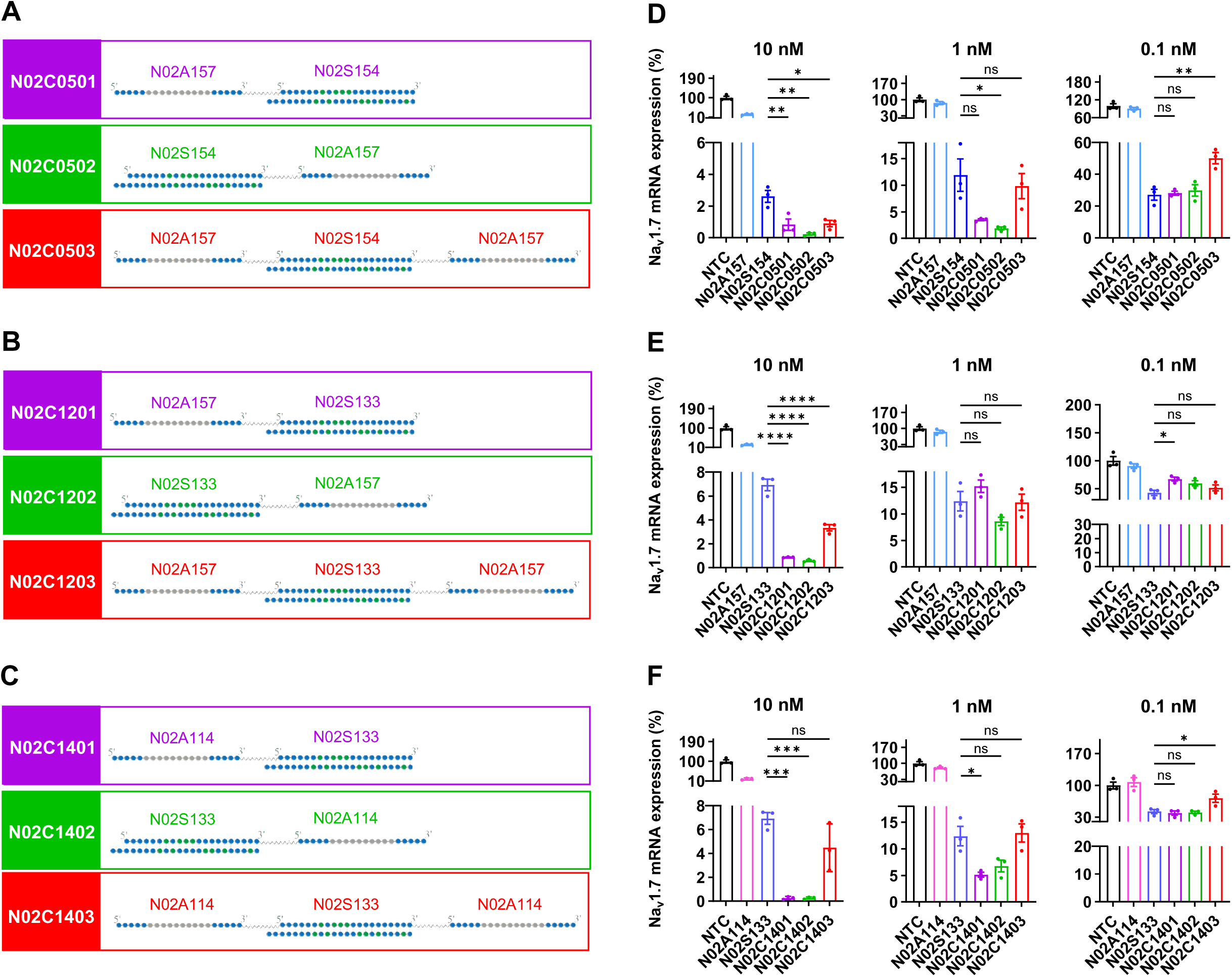
Characterization of nine additional ASO–siRNA conjugates. **(A-C)**, Schematic representation of conjugate architectures. Class 01 (purple): the 3’ end of the ASO is linked to the 5’ terminus of the siRNA sense strand (SS) via a PEG6 linker. Class 02 (green): the 5’ end of the ASO is linked to the 3’ terminus of the siRNA sense strand via a TEG6 linker. Class 03 (red): the siRNA SS is conjugated to two identical ASOs, with the 5’ terminus of the siRNA linked to the 3’ end of one ASO and the 3’ terminus linked to the 5’ end of the second ASO. (**D)** SK-N-AS cells were transfected with N02A157 (ASO), N02S154 (siRNA), or the corresponding conjugates at 0.1, 1, or 10 nM for 48 h. (**E)** SK-N-AS cells were transfected with N02A157 (ASO), N02S133 (siRNA), or the corresponding conjugates at 0.1, 1, or 10 nM for 48 h. (**F)** N02A114 (ASO), N02S133 (siRNA) and their corresponding conjugates were transfected into SK-N-AS cells at 0.1, 1 and 10 nM for 48h. Relative Nav1.7 mRNA levels were quantified by RT-qPCR. Statistical significance was determined using one-way ANOVA with Tukey’s multiple-comparisons test. n = 3 experimental replicates. Data are presented as mean ± SEM. ns *P* > 0.05, \**P* < 0.05, \*\**P* < 0.01, \*\*\**P* < 0.001, \*\*\*\**P* < 0.0001.

**Figure S4.**
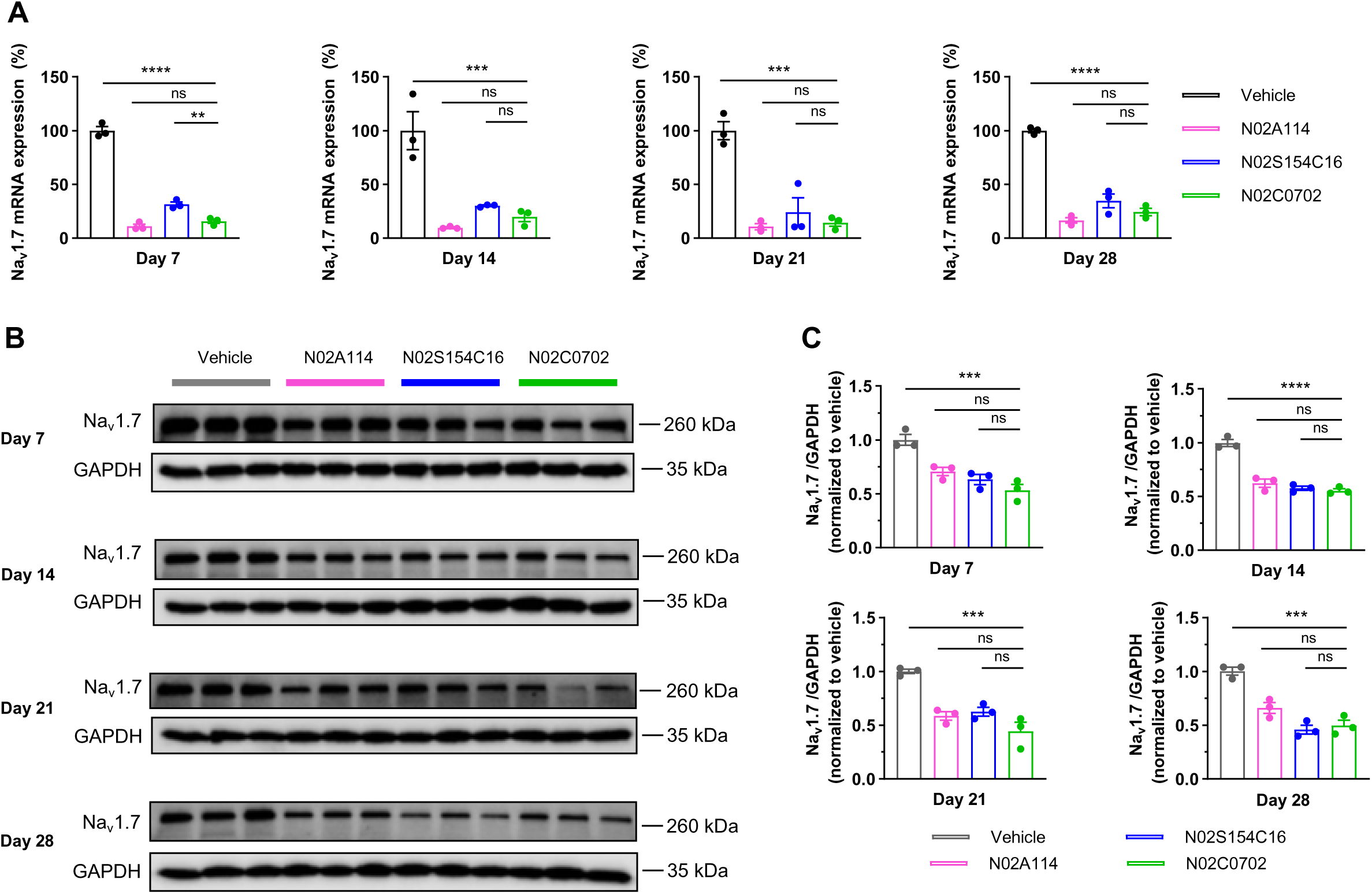
Comparison of the effects of N02C0702, N02A114, and N02S154C16 on Nav1.7 expression in spinal cord tissues. **(A, B)** Effects of intrathecal administration of N02A114 (1.5 mg /rat), N02S154C16 (3.0 mg /rat), and N02C0702 (4.5 mg /rat) on Nav1.7 mRNA (**A**) and protein (**B**) levels in DRG tissues at days 7, 14, 21, and 28 post-dose. n = 3 animals per group. **(C)** Quantification of Nav1.7 protein bands. Band intensities were measured using ImageJ software and normalized to GAPDH. n = 3 animals per group. Statistical significance was determined by one-way ANOVA with Dunnett’s multiple-comparisons test using N02C0702 as the reference group. Data are presented as mean ± SEM. ns *P* > 0.05, \**P* < 0.05, \*\**P* < 0.01, \*\*\**P* < 0.001, \*\*\*\**P* < 0.0001.

**Figure S5.**
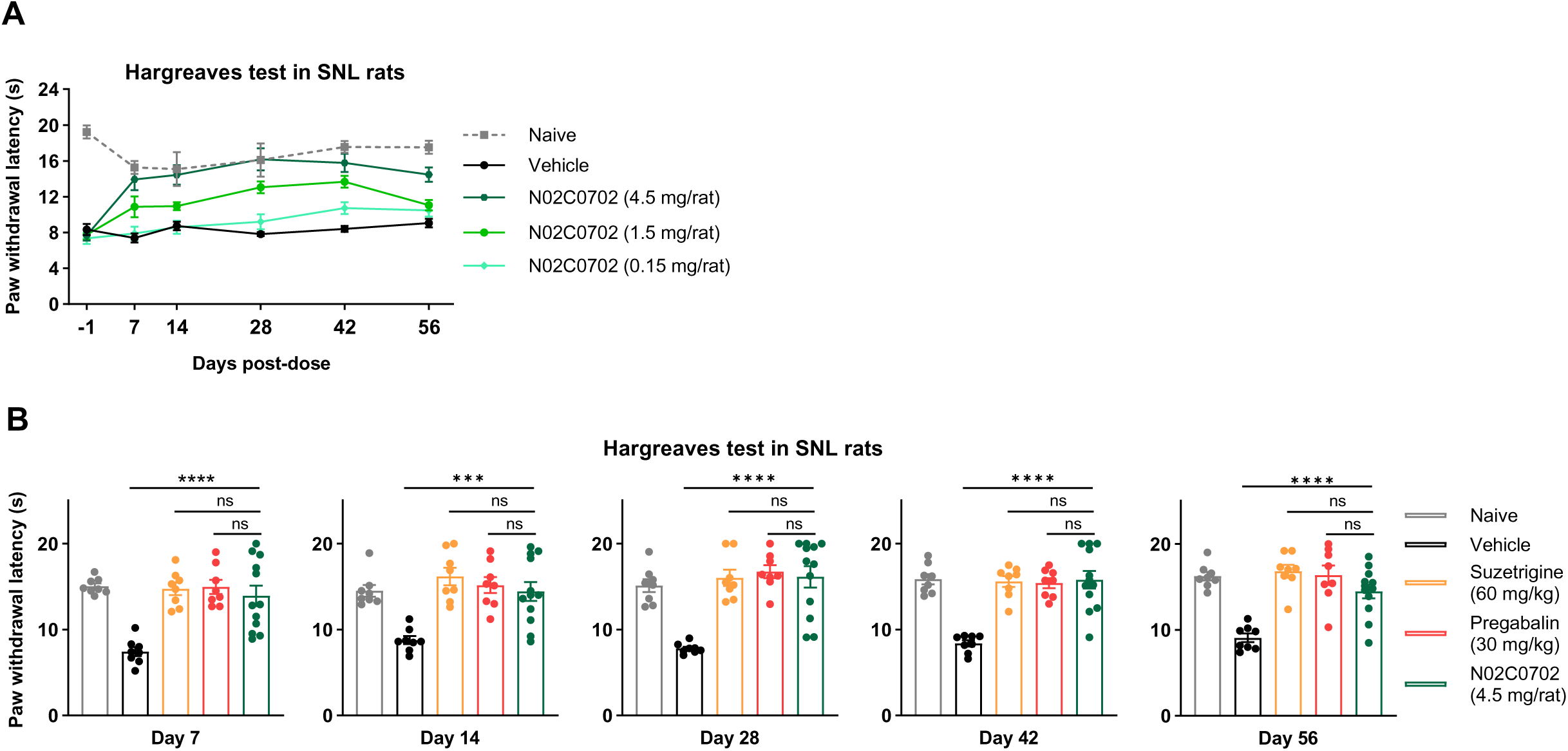
In vivo analgesic efficacy of N02C0702 in the Hargreaves test. One week after SNL surgery, rats received a single intrathecal injection of N02C0702 at 0.15, 1.5, or 4.5 mg /rat. Pregabalin (30 mg/kg) and suzetrigine (60 mg/kg) were administered orally 2 h prior to testing. Hargreaves tests were performed at the indicated time points post-dose. **(A)** Dose-dependent effects of N02C0702 on paw withdrawal latency (PWL) were measured at days 7, 14, 28, 42, and 56 post-SNL. \ **(B)** Comparison of PWLs among N02C0702 (4.5 mg/rat), suzetrigine (60 mg/kg), pregabalin (30 mg/kg), vehicle, and naive groups across time points. n = 8 animals per group for naive, vehicle, pregabalin, and suzetrigine; n = 12 animals for N02C0702. Statistical significance was determined by one-way ANOVA with Dunnett’s multiple-comparisons test using N02C0702 as the reference group. Data are presented as mean ± SEM. ns *P* > 0.05, \**P* < 0.05, \*\**P* < 0.01, \*\*\**P* < 0.001, \*\*\*\**P* < 0.0001.

## EXPERIMENTAL MODEL

### Oligonucleotides

The oligonucleotides used in this study were chemically synthesized and purified by RNase-free reversed-phase high-performance liquid chromatography (RP-HPLC) by WuXi TIDES (Shanghai, China).

Oligonucleotide synthesis was performed on a MerMade 12 synthesizer using standard solid-phase synthesis protocols. Gapmer antisense oligonucleotides (ASOs) were designed as 19–22-mer phosphorothioate-modified oligomers containing 5-mer 2’-O-methoxyethyl (2’-MOE) modifications at both the 5’ and 3’ terminal wings. During synthesis, 5-methylcytosine was incorporated in place of cytosine. siRNAs were modified using an advanced ESC design strategy. The two siRNA strands were synthesized separately and subsequently annealed to form duplexes, which were purified by size-exclusion chromatography to a purity of >90%. The conjugated sense strands were synthesized in the 3’→5’ direction, with a PEG6 linker introduced as a phosphoramidite monomer (DMTr-PEG6 amidite, CAS: 125607-09-2). Nusinersen (Cat. No. HY-112980) was purchased from MedChemExpress. The sequences and chemical modifications of all oligonucleotides are listed in **Tables S1, S2**.

### Cell culture

SK-N-AS cells (ATCC, CRL-2137), HeLa cells (ATCC, CCL-2; kindly provided by the Shanghai Institute of Biochemistry and Cell Biology), and Hepa1-6 cells (Cell Bank/Stem Cell Bank, Chinese Academy of Sciences, SCSP-512) were cultured in DMEM (Gibco, 11995040) supplemented with 10% fetal bovine serum (Gibco, 10091148), 1% penicillin–streptomycin (Gibco, 15140122), and 1% non-essential amino acids (NEAA; Gibco, 11140050). Hepa1-6 cells were additionally supplemented with 1% sodium pyruvate (Gibco, 11360070).

All animal procedures were conducted in accordance with the guidelines of the Institutional Animal Care and Use Committee (IACUC). Primary dorsal root ganglion (DRG) neurons were freshly isolated from 2–3-week-old Sprague–Dawley rats as previously described. Briefly, DRGs were harvested, minced with scissors, and incubated at 37 °C for 40 min in a digestion solution containing 0.25% trypsin–EDTA (Gibco, 25200072) and 0.2% collagenase type IV (Gibco, 17104019). After gentle trituration with pipettes, digestion was terminated by adding DMEM supplemented with 20% fetal bovine serum (FBS). The dissociated cells were centrifuged at 500 × g for 5 min, and the resulting pellet was resuspended in Primary Neuronal Cell Culture Medium (iCell, PriMed-iCel1-005). Cells were then plated onto 25 cm² flasks pre-coated with poly-D-lysine (Gibco, 14298806) to facilitate neuronal attachment. DRG neurons were maintained at 37 °C in a humidified incubator containing 5% CO₂ and 95% air for 2–3 days, after which they were collected and reseeded into 96-well plates for subsequent experiments.

### In vitro screening

SK-N-AS cells (1 × 10⁴ cells per well) or primary DRG cells (4 × 10⁴ cells per well) were seeded into 96-well plates and incubated for 24 h prior to transfection. Oligonucleotides were transfected using Lipofectamine RNAiMAX reagent (Invitrogen, 13778075) according to the manufacturer’s instructions. After 48 h of incubation, total RNA was extracted using a nucleic acid isolation reagent (DAAN Gene, #0621) with the KingFisher Flex Purification System (Thermo Fisher, A32681). RNA quantity and purity were assessed using a NanoDrop One spectrophotometer (Thermo Fisher, 840-317500), with acceptable A260/A280 ratios ranging from 1.8 to 2.1.

### Reverse transcription qPCR (RT-qPCR)

cDNA templates were generated by reverse transcription of total RNA using PrimeScript RT Master Mix (Takara, RR036B). Quantitative PCR (qPCR) was performed on a CFX384 Real-Time PCR System (Bio-Rad).

Target gene expression was quantified using TaqMan qPCR. Reactions were prepared using Premix Ex Taq (Takara, RR390B) according to the manufacturer’s instructions, with gene-specific primers and probes listed in **Table S4**.

Potential off-target gene expression was quantified using SYBR Green qPCR. RNA from SK-N-AS cells was prepared as described above. qPCR was performed in 384-well plates using PerfectStart Green qPCR SuperMix (TransGen, AQ601-01-V2) with gene-specific primers listed in **Table S4**.

Cycle threshold (Ct) values for each gene were normalized to the endogenous control GAPDH. Relative gene expression levels were calculated using the 2^−ΔΔCT method^81^.

### Human PBMC assay

Freshly isolated human PBMCs were purchased from Milecell Bio Co., Ltd. (Shanghai, China). The cells were seeded in 96-well plates (Corning, 3599) by 2 × 10^5^ cells per well, in 89% RPMI 1640 + 10% FBS + 1% penicillin/streptomycin. Cells were incubated at 37°C, 5% CO_2_ for 24 h. Next day, each ASO was serially diluted and added to 96-well plates at final nucleic acid concentration of 1.25, 0.31 and 0.078 μM, respectively. Cell supernatants were collected after 24 h incubation and stored at −80°C until processing for cytokine assay profiling. IL-6 (BioLegend, 430507) and TNF-α (ACROBiosystems, CRS-A002) levels were measured using sandwich ELISA kits according to the manufacturers’ instructions, and absorbance was read with a Varioskan LUX multimode microplate reader (Thermo Scientific, VL0L0TD0).

### psiCHECK assay

Off-target activity was assessed using the Dual-Glo® Luciferase Assay System (Promega, E2920). The psiCHECK™-2 vector contains a constitutively expressed firefly luciferase cassette and a multiple cloning site located in the 3’-UTR of the Renilla luciferase gene^82^. Oligonucleotides containing three tandem repeats of seed-matched (SM) sequences (**Table S5**) were synthesized with XhoI (New England Biolabs, R0146S) and NotI (New England Biolabs, R3189S) restriction sites at the ends. These tandem SM sequences were cloned into the 3’-UTR of the Renilla luciferase gene using the corresponding restriction enzymes. The resulting plasmids were used to evaluate seed region-dependent off-target activity of siRNAs. Firefly luciferase expressed from the psiCHECK™-2 vector served as an internal control for normalization. In the presence of off-target effects, siRNAs bind to the seed-matched sequences in the 3’-UTR of Renilla luciferase transcripts, leading to transcript degradation and reduced Renilla luciferase expression. Thus, increased off-target activity corresponds to lower Renilla luciferase activity.

HeLa cells were seeded at 4 × 10³ cells per well in white opaque 96-well plates. The following day, cells were co-transfected with siRNA (25, 6.25, 1.56, or 0.39 nM) and 80 ng of psiCHECK™-2-gSM reporter plasmid using 0.3 μL Lipofectamine 2000 (Invitrogen, 11668027) per well. Cells were harvested 48 h after transfection. Dual-Glo® Luciferase Reagent was first added to measure firefly luminescence, followed by the addition of Dual-Glo® Stop & Glo® Reagent to measure Renilla luminescence. Renilla luciferase activity was normalized to firefly luciferase activity for each well to control for transfection efficiency and ensure reproducible measurements.

### Caspase 3/7 assays

To assess Caspase 3/7 activity in cells, the Caspase-Glo® 3/7 Assay System (Promega, G8090) was added directly to the cells in a 1:1 volume ratio of reagent to sample. Following a 1.5-hour incubation at room temperature, luminescence was measured using a Varioskan LUX multimode microplate reader (Thermo Scientific, VL0L0TD0). Background luminescence from wells containing only culture medium was subtracted from the experimental readings. The relative Caspase 3/7 activity was then calculated as: 100% × (luminescence of treated sample) divided by (luminescence of vehicle control).

### Animals

Male Sprague-Dawley rats (about 200 g) were purchased from Vital River (Beijing, China) for *in vivo* analgesic evaluation. Male and female Sprague-Dawley rat pups (P15-P20) were purchased from the Vital River for primary DRG neuronal extraction and *in vitro* screening of oligonucleotides.

All experiments involving animals were performed in accordance with guidelines and regulations of the Institutional Animal Care and Use Committee (IACUC). The animal study protocol was approved by Shanghai Nyuen Biotech Ltd. Rats used in this study were housed in individually ventilated rat culture cages (Hongteng, 0251) in a dedicated, climate-controlled room maintained at 22°C and 45–50% humidity under a 12 h light/12 h dark cycle (lights on from 06:30 to 18:30). Animals were fed a Laboratory Rodent Diet (Labdiet, 5001) and had ad libitum access to reverse osmosis water.

### Rats IT dosing

Oligonucleotides were prepared at concentrations of up to 225 μg/mL for ASCs or 150 μg/mL for ASOs and siRNAs in artificial cerebrospinal fluid (aCSF) into 20 μL aliquots and IT injected in the dorsal spinal area, specifically between the L3 and L5 vertebrae, to adult male Sprague Dawley rats^83^. Prior to the surgical procedure, the rats were anesthetized with 2% isoflurane (Merck, 792632), then positioned on a heated mat (Stoelting, 53850R) and the injection site on the back was shaved and sanitized. A small incision was made to expose the spinal column, then the drug was injected using an insulin syringe. After the drug injection, gentle and sustained pressure was applied to the syringe plunger for approximately 30 seconds. The incision was then sutured, and the rats were placed supine on a warming pad for recovery.

### Tissue collection and RNA isolation

For sample collection, animals were euthanized by 2% isoflurane (Merck, 792632) and perfused with 150 mL of cold physiological saline through the left ventricle using a peristaltic pump. Cervical, thoracic, lumbar spinal cord and one side of lumbar DRG (L1-L6) were collected and stored in RNAlater stabilizing solution (Invitrogen, AM7021). All tissues were homogenized and lysed in TRIzol reagent (Invitrogen, 15596018CN), mixed with chloroform (Merck, 1.02445) and centrifuged to obtain an aqueous layer. Then, total RNA was isolated according to the RNeasy Mini Kit kit (QIAGEN, 74106) method. During isolation, 27U DNase I (TIANGEN, RT411) was used for in column DNA digestion.

### Western blotting

Western blotting was performed as previously described, with minor modifications^84^. The L4-L6 DRGs and the lumbar spinal cord were harvested from rats and immediately snap-frozen in liquid nitrogen. Total protein was extracted by lysis and homogenization in ice-cold NP-40 lysis buffer (Thermo Fisher, J60766. AK) containing Halt™ Protease Inhibitor Cocktail (Thermo Fisher Scientific, 87785). Following centrifugation at 16,000 × g for 10 min at 4°C, the supernatant was collected and protein concentrations were determined with a BCA kit (Thermo Fisher Scientific, 23227). After mixing the protein with 1× SDS-PAGE sample buffer and denaturing in 95℃ for 5 min, the samples were separated via electrophoresis on 10% polyacrylamide gels and transferred to 0.45 μm PVDF membranes (Thermo Fisher, 88585). The membranes were blocked with 5% non-fat milk (Santa Cruz, sc-2324) in 1×TBST for 1 h at room temperature, then incubated overnight at 4°C with gentle agitation in primary antibodies against Na_v_1.7 (1:200, Alomone Labs, ASC-008) and GAPDH (1:1000, Cell Signaling Technology, 2118S). A horseradish peroxidase (HRP)–conjugated goat anti-rabbit IgG antibody (1:1000, Cell Signaling Technology, 7074S) was applied to the membranes for 4 h at room temperature. The results were detected with an enhanced SuperSignal™ West Pico PLUS Chemiluminescent Substrate (Thermo Fisher, 34580) and the signal was visualized using a ChemiDoc MP Imaging System (Bio-Rad). The gray values of the western blot bands were quantified by ImageJ software.

### Spinal nerve ligation (SNL) Pain Model

The spinal nerve ligation (SNL) pain model was generated as previously described with minor modifications^63^. Animals were anesthetized by intraperitoneal injection with a mixture of xylazine (50 mg/kg) and ketamine (8 mg/kg). Ophthalmic ointment was applied to the rats’ eyes to prevent corneal dryness. The surgical area on the rats’ back was shaved, and the skin was disinfected three times with iodine tincture (Thermo Fisher, R260261) and 70% ethanol. A longitudinal incision was made posterior to the sacrum on the rat’s waist to expose the left paravertebral muscles. A retractor was used to separate the muscle tissue to expose the vertebrae. The left L5 and L6 spinal nerves were isolated and tightly ligated with 6-0 silk sutures (Fisher Scientific, NC9742105). The wound was sutured by Med Vet International Ethicon Vicryl Polyglactin 910 Plus Antibacterial Suture (Fisher Scientific, 50-283-0026). Postoperatively, animals were placed on a heating pad (Stoelting, 53850R), and returned to their cages once they had fully recovered and were able to move freely.

### CFA-induced inflammation pain model

The complete Freund’s adjuvant (CFA)-induced inflammation pain model was generated as previously described with minor modifications^85^. After one week of adaptive feeding, rats received a subcutaneous injection of 100 μL CFA (Merck, AR001) at a concentration of 1 mg/mL into the plantar center of the left hind paw to induce inflammatory pain. Mechanical hyperalgesia was assessed 24 h after CFA induction to confirm the success of inflammatory pain model.

### Measurement of mechanical allodynia (von Frey testing)

The von Frey testing of rats was generated as previously described with minor modifications^86^. Each rat was placed in a transparent cubicle with a wire mesh floor and allowed to acclimate to the apparatus for 1 h. Mechanical sensitivity was assessed using a series of calibrated von Frey filaments (0.4-15.1 g; Stoelting Co., USA, 57814) applied to the plantar surface of the hind paw. Each filament was applied vertically for 3-4 s with sufficient force to produce slight bending. Each filament was tested five times per trial. A brisk paw withdrawal was considered a positive response, and a consistent response in at least three of five applications was defined as the threshold response. The up–down method described by Dixon was used to convert the pattern of positive and negative responses into a 50% withdrawal threshold value.

### Measurement of thermal hyperalgesia (Hargreaves test)

Rats were assessed for thermal hyperalgesia as described in previous study^5^. Hargreaves test was performed with Hot plate (BioMed, SA702). Rats were brought into the testing room and allowed to acclimate for at least 30 min prior to testing. The hot plate surface was cleaned with an appropriate disinfectant before each trial. The plate was heated to 55°C, as monitored by the built-in digital thermometer, with temperature maintained within ±0.2°C. Each animal was placed on the heated surface inside a transparent cylinder, and the timer was initiated immediately. The latency to the first nociceptive response—defined as hind paw licking, hind paw flicking, or jumping—was recorded, and the animal was promptly removed upon exhibiting this response. If no response occurred within 20 s at the maximum temperature, the trial was terminated and the animal was removed to prevent thermal injury. Each rat was tested three times to obtain an average reaction time. The hot plate surface was disinfected between testing of individual animals.

### RNA sequencing

SK-N-AS cells (ATCC, CRL-2137) were cultured in 24-well plates and transfected with the indicated oligonucleotides using Lipofectamine RNAiMAX reagent (Invitrogen, 13778075) for 24 h. Total RNA was extracted using RNeasy Mini Kits (Qiagen), and the integrity of the RNA was assessed with the Agilent 2100 Bioanalyzer. Libraries were prepared using the azyme-NR606-02 VAHTS® Universal library preparation kit for sequencing on an Illumina NovaSeq 6000 platform (Illumina Inc., San Diego, CA, USA).

Sequencing data were mapped to the GRCh38 reference genome using STAR software, followed by quantification with HTSeq-Count. Differential expression analysis was conducted using the DESeq2 package, and log2 Fold Change values were adjusted with lfcShrink^87^. Genes were selected as differentially expressed if they had an absolute value of log2 Fold Change greater than 1 and a Benjamini-Hochberg (BH) corrected p-value of less than 0.05. Graphs were generated using the R package ggplot2.

### Data analysis

Statistical analyses were performed using GraphPad PRISM software (version 10.1.2). The specific statistical test used in each experiment was depicted in the legends of corresponding figures. Results are presented as mean ± standard error of mean (SEM); error bars represent SEMs. Statistical significance is shown by asterisks, \**P* < 0.05, \*\**P* < 0.01, \*\*\**P* < 0.001 and \*\*\*\**P* < 0.0001. Differences among groups were analyzed using one-way or two-way analysis of variance (ANOVA), as appropriate.

## RESOURCE AVAILABILITY

### Lead contact

Requests for further information, resources, and reagents should be directed to and will be fulfilled by the lead contact, Bin Ren and Guohao Wang.

### Materials availability

This study generates new, unique ASCs reagents that can be request from the Dr. Bin Ren and Dr. Guohao Wang.

## REFERENCES

1. Cohen, S.P., Vase, L., and Hooten, W.M. (2021). Chronic pain: an update on burden, best practices, and new advances. Lancet 397, 2082–2097. 10.1016/S0140-6736(21)00393-7.

2. Stubhaug, A., Hansen, J.L., Hallberg, S., Gustavsson, A., Eggen, A.E., and Nielsen, C.S. (2024). The costs of chronic pain-Long-term estimates. Eur J Pain 28, 960–977. 10.1002/ejp.2234.

3. Kang, Y., Trewern, L., Jackman, J., McCartney, D., and Soni, A. (2023). Chronic pain: definitions and diagnosis. BMJ 381, e076036. 10.1136/bmj-2023-076036.

4. Volkow, N.D., and McLellan, A.T. (2016). Opioid Abuse in Chronic Pain--Misconceptions and Mitigation Strategies. N Engl J Med 374, 1253–1263. 10.1056/NEJMra1507771.

5. Hargreaves, K., Dubner, R., Brown, F., Flores, C., and Joris, J. (1988). A new and sensitive method for measuring thermal nociception in cutaneous hyperalgesia. Pain 32, 77–88. 10.1016/0304-3959(88)90026-7.

6. Hauser, W., Morlion, B., Vowles, K.E., Bannister, K., Buchser, E., Casale, R., Chenot, J.F., Chumbley, G., Drewes, A.M., Dom, G., et al. (2021). European* clinical practice recommendations on opioids for chronic noncancer pain - Part 1: Role of opioids in the management of chronic noncancer pain. Eur J Pain 25, 949–968. 10.1002/ejp.1736.

7. Jayakar, S., Shim, J., Jo, S., Bean, B.P., Singec, I., and Woolf, C.J. (2021). Developing nociceptor-selective treatments for acute and chronic pain. Sci Transl Med 13, eabj9837. 10.1126/scitranslmed.abj9837.

8. Papapetropoulos, A., Topouzis, S., Alexander, S.P.H., Cortese-Krott, M.M., Helyes, Z., Martemyanov, K., Mauro, C., Nagercoil, N., Panettieri, R.A., Jr., Patel, H.H., et al. (2026). Novel drugs approved by the EMA, the FDA and the MHRA in 2025: A year in review. Br J Pharmacol. 10.1111/bph.70376.

9. Jones, J., Correll, D.J., Lechner, S.M., Jazic, I., Miao, X., Shaw, D., Simard, C., Osteen, J.D., Hare, B., Beaton, A., et al. (2023). Selective Inhibition of Na(V)1.8 with VX-548 for Acute Pain. N Engl J Med 389, 393–405. 10.1056/NEJMoa2209870.

10. Bertoch, T., D’Aunno, D., McCoun, J., Solanki, D., Taber, L., Urban, J., Oswald, J., Swisher, M.W., Tian, S., Miao, X., et al. (2025). Suzetrigine, a Nonopioid Na V 1.8 Inhibitor for Treatment of Moderate-to-severe Acute Pain: Two Phase 3 Randomized Clinical Trials. Anesthesiology 142, 1085–1099. 10.1097/ALN.0000000000005460.

11. Kingwell, K. (2025). Na(V)1.8 inhibitor poised to provide opioid-free pain relief. Nat Rev Drug Discov 24, 3–5. 10.1038/d41573-024-00203-3.

12. Khan, A., Irshad, M., Javed, Z., Naz, Z., and Fatima, M. (2025). Beyond opioids: FDA-approved suzetrigine offers hope for acute pain management. Ann Med Surg (Lond) 87, 4023–4025. 10.1097/MS9.0000000000003462.

13. Bennett, D.L., Clark, A.J., Huang, J., Waxman, S.G., and Dib-Hajj, S.D. (2019). The Role of Voltage-Gated Sodium Channels in Pain Signaling. Physiol Rev 99, 1079–1151. 10.1152/physrev.00052.2017.

14. Goodwin, G., and McMahon, S.B. (2021). The physiological function of different voltage-gated sodium channels in pain. Nat Rev Neurosci 22, 263–274. 10.1038/s41583-021-00444-w.

15. Waxman, S.G. (2023). Targeting a Peripheral Sodium Channel to Treat Pain. N Engl J Med 389, 466–469. 10.1056/NEJMe2305708.

16. Cox, J.J., Reimann, F., Nicholas, A.K., Thornton, G., Roberts, E., Springell, K., Karbani, G., Jafri, H., Mannan, J., Raashid, Y., et al. (2006). An SCN9A channelopathy causes congenital inability to experience pain. Nature 444, 894–898. 10.1038/nature05413.

17. Cox, J.J., Sheynin, J., Shorer, Z., Reimann, F., Nicholas, A.K., Zubovic, L., Baralle, M., Wraige, E., Manor, E., Levy, J., et al. (2010). Congenital insensitivity to pain: novel SCN9A missense and in-frame deletion mutations. Hum Mutat 31, E1670–1686. 10.1002/humu.21325.

18. Nilsen, K.B., Nicholas, A.K., Woods, C.G., Mellgren, S.I., Nebuchennykh, M., and Aasly, J. (2009). Two novel SCN9A mutations causing insensitivity to pain. Pain 143, 155–158. 10.1016/j.pain.2009.02.016.

19. Yang, Y., Wang, Y., Li, S., Xu, Z., Li, H., Ma, L., Fan, J., Bu, D., Liu, B., Fan, Z., et al. (2004). Mutations in SCN9A, encoding a sodium channel alpha subunit, in patients with primary erythermalgia. J Med Genet 41, 171–174. 10.1136/jmg.2003.012153.

20. Dib-Hajj, S.D., Rush, A.M., Cummins, T.R., Hisama, F.M., Novella, S., Tyrrell, L., Marshall, L., and Waxman, S.G. (2005). Gain-of-function mutation in Nav1.7 in familial erythromelalgia induces bursting of sensory neurons. Brain 128, 1847–1854. 10.1093/brain/awh514.

21. Fertleman, C.R., Baker, M.D., Parker, K.A., Moffatt, S., Elmslie, F.V., Abrahamsen, B., Ostman, J., Klugbauer, N., Wood, J.N., Gardiner, R.M., and Rees, M. (2006). SCN9A mutations in paroxysmal extreme pain disorder: allelic variants underlie distinct channel defects and phenotypes. Neuron 52, 767–774. 10.1016/j.neuron.2006.10.006.

22. Faber, C.G., Hoeijmakers, J.G., Ahn, H.S., Cheng, X., Han, C., Choi, J.S., Estacion, M., Lauria, G., Vanhoutte, E.K., Gerrits, M.M., et al. (2012). Gain of function Nanu1.7 mutations in idiopathic small fiber neuropathy. Ann Neurol 71, 26–39. 10.1002/ana.22485.

23. Vetter, I., Deuis, J.R., Mueller, A., Israel, M.R., Starobova, H., Zhang, A., Rash, L.D., and Mobli, M. (2017). Na(V)1.7 as a pain target - From gene to pharmacology. Pharmacol Ther 172, 73–100. 10.1016/j.pharmthera.2016.11.015.

24. Alsaloum, M., Higerd, G.P., Effraim, P.R., and Waxman, S.G. (2020). Status of peripheral sodium channel blockers for non-addictive pain treatment. Nat Rev Neurol 16, 689–705. 10.1038/s41582-020-00415-2.

25. Alsaloum, M., Dib-Hajj, S.D., Page, D.A., Ruben, P.C., Krainer, A.R., and Waxman, S.G. (2025). Voltage-gated sodium channels in excitable cells as drug targets. Nat Rev Drug Discov 24, 358–378. 10.1038/s41573-024-01108-x.

26. Kotecha, M., Cheshire, W.P., Finnigan, H., Giblin, K., Naik, H., Palmer, J., Tate, S., and Zakrzewska, J.M. (2020). Design of Phase 3 Studies Evaluating Vixotrigine for Treatment of Trigeminal Neuralgia. J Pain Res 13, 1601–1609. 10.2147/JPR.S247182.

27. Zakrzewska, J.M., Palmer, J., Morisset, V., Giblin, G.M., Obermann, M., Ettlin, D.A., Cruccu, G., Bendtsen, L., Estacion, M., Derjean, D., et al. (2017). Safety and efficacy of a Nav1.7 selective sodium channel blocker in patients with trigeminal neuralgia: a double-blind, placebo-controlled, randomised withdrawal phase 2a trial. Lancet Neurol 16, 291–300. 10.1016/S1474-4422(17)30005-4.

28. McDonnell, A., Collins, S., Ali, Z., Iavarone, L., Surujbally, R., Kirby, S., and Butt, R.P. (2018). Efficacy of the Nav1.7 blocker PF-05089771 in a randomised, placebo-controlled, double-blind clinical study in subjects with painful diabetic peripheral neuropathy. Pain 159, 1465–1476. 10.1097/j.pain.0000000000001227.

29. Eagles, D.A., Chow, C.Y., and King, G.F. (2022). Fifteen years of Na(V) 1.7 channels as an analgesic target: Why has excellent in vitro pharmacology not translated into in vivo analgesic efficacy? Br J Pharmacol 179, 3592–3611. 10.1111/bph.15327.

30. Clare, J.J., Tate, S.N., Nobbs, M., and Romanos, M.A. (2000). Voltage-gated sodium channels as therapeutic targets. Drug Discov Today 5, 506–520. 10.1016/s1359-6446(00)01570-1.

31. Loussouarn, G., Sternberg, D., Nicole, S., Marionneau, C., Le Bouffant, F., Toumaniantz, G., Barc, J., Malak, O.A., Fressart, V., Pereon, Y., et al. (2015). Physiological and Pathophysiological Insights of Nav1.4 and Nav1.5 Comparison. Front Pharmacol 6, 314. 10.3389/fphar.2015.00314.

32. Bennett, D.L., and Woods, C.G. (2014). Painful and painless channelopathies. Lancet Neurol 13, 587–599. 10.1016/S1474-4422(14)70024-9.

33. Lischka, A., Lassuthova, P., Cakar, A., Record, C.J., Van Lent, J., Baets, J., Dohrn, M.F., Senderek, J., Lampert, A., Bennett, D.L., et al. (2022). Genetic pain loss disorders. Nat Rev Dis Primers 8, 41. 10.1038/s41572-022-00365-7.

34. Jadhav, V., Vaishnaw, A., Fitzgerald, K., and Maier, M.A. (2024). RNA interference in the era of nucleic acid therapeutics. Nat Biotechnol 42, 394–405. 10.1038/s41587-023-02105-y.

35. Hu, B., Zhong, L., Weng, Y., Peng, L., Huang, Y., Zhao, Y., and Liang, X.J. (2020). Therapeutic siRNA: state of the art. Signal Transduct Target Ther 5, 101. 10.1038/s41392-020-0207-x.

36. Tang, Q., and Khvorova, A. (2024). RNAi-based drug design: considerations and future directions. Nat Rev Drug Discov 23, 341–364. 10.1038/s41573-024-00912-9.

37. Crooke, S.T., Witztum, J.L., Bennett, C.F., and Baker, B.F. (2018). RNA-Targeted Therapeutics. Cell Metab 27, 714–739. 10.1016/j.cmet.2018.03.004.

38. Roberts, T.C., Langer, R., and Wood, M.J.A. (2020). Advances in oligonucleotide drug delivery. Nat Rev Drug Discov 19, 673–694. 10.1038/s41573-020-0075-7.

39. Androsavich, J.R. (2024). Frameworks for transformational breakthroughs in RNA-based medicines. Nat Rev Drug Discov 23, 421–444. 10.1038/s41573-024-00943-2.

40. Crooke, S.T., Baker, B.F., Crooke, R.M., and Liang, X.H. (2021). Antisense technology: an overview and prospectus. Nat Rev Drug Discov 20, 427–453. 10.1038/s41573-021-00162-z.

41. Sun, X., Setrerrahmane, S., Li, C., Hu, J., and Xu, H. (2024). Nucleic acid drugs: recent progress and future perspectives. Signal Transduct Target Ther 9, 316. 10.1038/s41392-024-02035-4.

42. Anand, P., Zhang, Y., Patil, S., and Kaur, K. (2025). Metabolic Stability and Targeted Delivery of Oligonucleotides: Advancing RNA Therapeutics Beyond The Liver. J Med Chem 68, 6870–6896. 10.1021/acs.jmedchem.4c02528.

43. Guo, S., Zhang, M., and Huang, Y. (2024). Three ‘E’ challenges for siRNA drug development. Trends Mol Med 30, 13–24. 10.1016/j.molmed.2023.10.005.

44. Kulkarni, J.A., Witzigmann, D., Thomson, S.B., Chen, S., Leavitt, B.R., Cullis, P.R., and van der Meel, R. (2021). The current landscape of nucleic acid therapeutics. Nat Nanotechnol 16, 630–643. 10.1038/s41565-021-00898-0.

45. Crnkovic-Mertens, I., Semzow, J., Hoppe-Seyler, F., and Butz, K. (2006). Isoform-specific silencing of the Livin gene by RNA interference defines Livin beta as key mediator of apoptosis inhibition in HeLa cells. J Mol Med (Berl) 84, 232–240. 10.1007/s00109-005-0021-5.

46. Ostergaard, M.E., Southwell, A.L., Kordasiewicz, H., Watt, A.T., Skotte, N.H., Doty, C.N., Vaid, K., Villanueva, E.B., Swayze, E.E., Bennett, C.F., et al. (2013). Rational design of antisense oligonucleotides targeting single nucleotide polymorphisms for potent and allele selective suppression of mutant Huntingtin in the CNS. Nucleic Acids Res 41, 9634–9650. 10.1093/nar/gkt725.

47. Conroy, F., Miller, R., Alterman, J.F., Hassler, M.R., Echeverria, D., Godinho, B., Knox, E.G., Sapp, E., Sousa, J., Yamada, K., et al. (2022). Chemical engineering of therapeutic siRNAs for allele-specific gene silencing in Huntington’s disease models. Nat Commun 13, 5802. 10.1038/s41467-022-33061-x.

48. Egli, M., and Manoharan, M. (2023). Chemistry, structure and function of approved oligonucleotide therapeutics. Nucleic Acids Res 51, 2529–2573. 10.1093/nar/gkad067.

49. Khvorova, A., and Watts, J.K. (2017). The chemical evolution of oligonucleotide therapies of clinical utility. Nat Biotechnol 35, 238–248. 10.1038/nbt.3765.

50. Wang, S., Weissman, D., and Dong, Y. (2025). RNA chemistry and therapeutics. Nat Rev Drug Discov 24, 828–851. 10.1038/s41573-025-01237-x.

51. Ray, K.K., Wright, R.S., Kallend, D., Koenig, W., Leiter, L.A., Raal, F.J., Bisch, J.A., Richardson, T., Jaros, M., Wijngaard, P.L.J., et al. (2020). Two Phase 3 Trials of Inclisiran in Patients with Elevated LDL Cholesterol. N Engl J Med 382, 1507–1519. 10.1056/NEJMoa1912387.

52. Raal, F.J., Kallend, D., Ray, K.K., Turner, T., Koenig, W., Wright, R.S., Wijngaard, P.L.J., Curcio, D., Jaros, M.J., Leiter, L.A., et al. (2020). Inclisiran for the Treatment of Heterozygous Familial Hypercholesterolemia. N Engl J Med 382, 1520–1530. 10.1056/NEJMoa1913805.

53. JOURNAVX (2025) JOURNAVX (suzetrigine) tablets, for oral use.

54. LYRICA (2018) LYRICA (pregabalin) Capsules, CV-accessdata.

55. Kole, R., Krainer, A.R., and Altman, S. (2012). RNA therapeutics: beyond RNA interference and antisense oligonucleotides. Nat Rev Drug Discov 11, 125–140. 10.1038/nrd3625.

56. Pollak, A.J., Zhao, L., and Crooke, S.T. (2023). Systematic Analysis of Chemical Modifications of Phosphorothioate Antisense Oligonucleotides that Modulate Their Innate Immune Response. Nucleic Acid Ther 33, 95–107. 10.1089/nat.2022.0067.

57. Burel, S.A., Machemer, T., Baker, B.F., Kwoh, T.J., Paz, S., Younis, H., and Henry, S.P. (2022). Early-Stage Identification and Avoidance of Antisense Oligonucleotides Causing Species-Specific Inflammatory Responses in Human Volunteer Peripheral Blood Mononuclear Cells. Nucleic Acid Ther 32, 457–472. 10.1089/nat.2022.0033.

58. Klinman, D.M. (2004). Immunotherapeutic uses of CpG oligodeoxynucleotides. Nat Rev Immunol 4, 249–258. 10.1038/nri1329.

59. Dieckmann, A., Hagedorn, P.H., Burki, Y., Brugmann, C., Berrera, M., Ebeling, M., Singer, T., and Schuler, F. (2018). A Sensitive In Vitro Approach to Assess the Hybridization-Dependent Toxic Potential of High Affinity Gapmer Oligonucleotides. Mol Ther Nucleic Acids 10, 45–54. 10.1016/j.omtn.2017.11.004.

60. Shen, W., De Hoyos, C.L., Migawa, M.T., Vickers, T.A., Sun, H., Low, A., Bell, T.A., 3rd, Rahdar, M., Mukhopadhyay, S., Hart, C.E., et al. (2019). Chemical modification of PS-ASO therapeutics reduces cellular protein-binding and improves the therapeutic index. Nat Biotechnol 37, 640–650. 10.1038/s41587-019-0106-2.

61. Jackson, A.L., and Linsley, P.S. (2010). Recognizing and avoiding siRNA off-target effects for target identification and therapeutic application. Nat Rev Drug Discov 9, 57–67. 10.1038/nrd3010.

62. Ui-Tei, K., Naito, Y., Nishi, K., Juni, A., and Saigo, K. (2008). Thermodynamic stability and Watson-Crick base pairing in the seed duplex are major determinants of the efficiency of the siRNA-based off-target effect. Nucleic Acids Res 36, 7100–7109. 10.1093/nar/gkn902.

63. Chung, J.M., Kim, H.K., and Chung, K. (2004). Segmental spinal nerve ligation model of neuropathic pain. Methods Mol Med 99, 35–45. 10.1385/1-59259-770-X:035.

64. Brown, K.M., Nair, J.K., Janas, M.M., Anglero-Rodriguez, Y.I., Dang, L.T.H., Peng, H., Theile, C.S., Castellanos-Rizaldos, E., Brown, C., Foster, D., et al. (2022). Expanding RNAi therapeutics to extrahepatic tissues with lipophilic conjugates. Nat Biotechnol 40, 1500–1508. 10.1038/s41587-022-01334-x.

65. Raymond, C.K., Castle, J., Garrett-Engele, P., Armour, C.D., Kan, Z., Tsinoremas, N., and Johnson, J.M. (2004). Expression of alternatively spliced sodium channel alpha-subunit genes. Unique splicing patterns are observed in dorsal root ganglia. J Biol Chem 279, 46234–46241. 10.1074/jbc.M406387200.

66. Yang, J., Xie, Y.F., Smith, R., Ratte, S., and Prescott, S.A. (2025). Discordance between preclinical and clinical testing of Na V 1.7-selective inhibitors for pain. Pain 166, 481–501. 10.1097/j.pain.0000000000003425.

67. Wu, H., Lima, W.F., Zhang, H., Fan, A., Sun, H., and Crooke, S.T. (2004). Determination of the role of the human RNase H1 in the pharmacology of DNA-like antisense drugs. J Biol Chem 279, 17181–17189. 10.1074/jbc.M311683200.

68. Crooke, S.T. (2017). Molecular Mechanisms of Antisense Oligonucleotides. Nucleic Acid Ther 27, 70–77. 10.1089/nat.2016.0656.

69. Liu, J., Carmell, M.A., Rivas, F.V., Marsden, C.G., Thomson, J.M., Song, J.J., Hammond, S.M., Joshua-Tor, L., and Hannon, G.J. (2004). Argonaute2 is the catalytic engine of mammalian RNAi. Science 305, 1437–1441. 10.1126/science.1102513.

70. Roberts, T.C. (2015). The microRNA Machinery. Adv Exp Med Biol 887, 15–30. 10.1007/978-3-319-22380-3_2.

71. Schurmann, N., Trabuco, L.G., Bender, C., Russell, R.B., and Grimm, D. (2013). Molecular dissection of human Argonaute proteins by DNA shuffling. Nat Struct Mol Biol 20, 818–826. 10.1038/nsmb.2607.

72. Sarli, S.L., Fakih, H.H., Kelly, K., Devi, G., Rembetsy-Brown, J.M., McEachern, H.R., Ferguson, C.M., Echeverria, D., Lee, J., Sousa, J., et al. (2024). Quantifying the activity profile of ASO and siRNA conjugates in glioblastoma xenograft tumors in vivo. Nucleic Acids Res 52, 4799–4817. 10.1093/nar/gkae260.

73. Kang, M., Lin, W.H., Li, Y., Xu, F., Zhou, X., Si, C., Dai, J., He, J., Schacht, I., Gan, Z., et al. (2025). SCAD: A modular platform for efficient delivery of duplex RNA to the CNS and beyond. Mol Ther Nucleic Acids 36, 102757. 10.1016/j.omtn.2025.102757.

74. Shields, S.D., Deng, L., Reese, R.M., Dourado, M., Tao, J., Foreman, O., Chang, J.H., and Hackos, D.H. (2018). Insensitivity to Pain upon Adult-Onset Deletion of Nav1.7 or Its Blockade with Selective Inhibitors. J Neurosci 38, 10180–10201. 10.1523/JNEUROSCI.1049-18.2018.

75. Miller, T.M., Cudkowicz, M.E., Shaw, P.J., Genge, A., Sobue, G., Bucelli, R.C., Chio, A., Van Damme, P., Ludolph, A.C., Glass, J.D., et al. (2026). Long-Term Tofersen in SOD1 Amyotrophic Lateral Sclerosis. JAMA Neurol 83, 115–125. 10.1001/jamaneurol.2025.4946.

76. Mercuri, E., Darras, B.T., Chiriboga, C.A., Day, J.W., Campbell, C., Connolly, A.M., Iannaccone, S.T., Kirschner, J., Kuntz, N.L., Saito, K., et al. (2018). Nusinersen versus Sham Control in Later-Onset Spinal Muscular Atrophy. N Engl J Med 378, 625–635. 10.1056/NEJMoa1710504.

77. Xiao, B., Wang, S., Pan, Y., Zhi, W., Gu, C., Guo, T., Zhai, J., Li, C., Chen, Y.Q., and Wang, R. (2025). Development, opportunities, and challenges of siRNA nucleic acid drugs. Mol Ther Nucleic Acids 36, 102437. 10.1016/j.omtn.2024.102437.

78. Study of ARO-DIMERPA in Adult Participants With Mixed Hyperlipidemia. (2025). https://clinicaltrials.gov/study/NCT07223658.

79. A Study to Evaluate COR-1004 in Adult Volunteers. (2025). https://clinicaltrials.gov/study/NCT07229118.

80. Alterman, J.F., Godinho, B., Hassler, M.R., Ferguson, C.M., Echeverria, D., Sapp, E., Haraszti, R.A., Coles, A.H., Conroy, F., Miller, R., et al. (2019). A divalent siRNA chemical scaffold for potent and sustained modulation of gene expression throughout the central nervous system. Nat Biotechnol 37, 884–894. 10.1038/s41587-019-0205-0.

81. Yin, W., Rajvanshi, P.K., Rogers, H.M., Yoshida, T., Kopp, J.B., An, X., Gassmann, M., and Noguchi, C.T. (2024). Erythropoietin regulates energy metabolism through EPO-EpoR-RUNX1 axis. Nat Commun 15, 8114. 10.1038/s41467-024-52352-z.

82. Zhou, J., Song, M.S., Jacobi, A.M., Behlke, M.A., Wu, X., and Rossi, J.J. (2012). Deep Sequencing Analyses of DsiRNAs Reveal the Influence of 3’ Terminal Overhangs on Dicing Polarity, Strand Selectivity, and RNA Editing of siRNAs. Mol Ther Nucleic Acids 1, e17. 10.1038/mtna.2012.6.

83. Fischer, G., Kostic, S., Nakai, H., Park, F., Sapunar, D., Yu, H., and Hogan, Q. (2011). Direct injection into the dorsal root ganglion: technical, behavioral, and histological observations. J Neurosci Methods 199, 43–55. 10.1016/j.jneumeth.2011.04.021.

84. Wang, G., Peng, S., Reyes Mendez, M., Keramidas, A., Castellano, D., Wu, K., Han, W., Tian, Q., Dong, L., Li, Y., and Lu, W. (2024). The TMEM132B-GABA(A) receptor complex controls alcohol actions in the brain. Cell 187, 6649–6668 e6635. 10.1016/j.cell.2024.09.006.

85. Zollner, C., Mousa, S.A., Fischer, O., Rittner, H.L., Shaqura, M., Brack, A., Shakibaei, M., Binder, W., Urban, F., Stein, C., and Schafer, M. (2008). Chronic morphine use does not induce peripheral tolerance in a rat model of inflammatory pain. J Clin Invest 118, 1065–1073. 10.1172/JCI25911.

86. Pertin, M., Allchorne, A.J., Beggah, A.T., Woolf, C.J., and Decosterd, I. (2007). Delayed sympathetic dependence in the spared nerve injury (SNI) model of neuropathic pain. Mol Pain 3, 21. 10.1186/1744-8069-3-21.

87. Mann, A.E., Chakraborty, B., O’Connell, L.M., Nascimento, M.M., Burne, R.A., and Richards, V.P. (2024). Heterogeneous lineage-specific arginine deiminase expression within dental microbiome species. Microbiol Spectr 12, e0144523. 10.1128/spectrum.01445-23.

