## Supplementary material for "Sustained Relief of Chronic Pain via a Nav1.7-Targeting ASO–siRNA Conjugate": Supplental Table1

### Table S1: The sequences and modifications of ASOs

| ASO# | modified_sequence | Length |
|---|---|---|
| N02A003 | moeCMe*moeA*moeT*moeCMe*moeA*dT*dG*dT*dA*dA*dA*dCMe*dA*dA*dT*dA*dA*moeG*moeG*moeCMe*moeA*moeCMe | 22 |
| N02A005 | moeG*moeT*moeG*moeT*moeT*dT*dA*dA*dA*dA*dCMe*dT*dA*dT*dG*dCMe*dA*moeA*moeA*moeT*moeG*moeG | 22 |
| N02A025 | moeT*moeT*moeG*moeCMe*moeA*dA*dG*dA*dT*dCMe*dT*dA*dCMe*dA*dA*dA*dA*moeG*moeG*moeA*moeT*moeCMe | 22 |
| N02A026 | moeA*moeG*moeCMe*moeCMe*moeT*dCMe*dT*dT*dG*dCMe*dA*dA*dG*dG*dA*moeT*moeT*moeT*moeT*moeT | 20 |
| N02A027 | moeCMe*moeT*moeT*moeT*moeCMe*dT*dT*dCMe*dCMe*dA*dT*dCMe*dT*dG*dT*dG*moeA*moeA*moeCMe*moeG*moeA | 21 |
| N02A040 | moeCMe*moeT*moeT*moeT*moeCMe*dA*dG*dG*dT*dT*dT*dT*dT*dCMe*dCMe*dA*dT*moeCMe*moeG*moeCMe*moeA*moeCMe | 22 |
| N02A046 | moeT*moeA*moeT*moeG*moeCMe*dA*dA*dA*dT*dG*dG*dT*dA*dA*dT*dT*dG*moeCMe*moeA*moeA*moeG*moeA | 22 |
| N02A048 | moeG*moeT*moeT*moeG*moeCMe*dCMe*dT*dT*dA*dT*dCMe*dT*dA*dT*dT*dA*moeT*moeCMe*moeA*moeCMe*moeCMe | 21 |
| N02A050 | moeA*moeCMe*moeT*moeT*moeT*dCMe*dA*dG*dG*dT*dT*dT*dT*dT*dCMe*dCMe*moeA*moeT*moeCMe*moeG*moeCMe | 21 |
| N02A051 | moeA*moeT*moeCMe*moeCMe*moeA*dA*dT*dA*dT*dG*dG*dA*dG*dA*dG*moeCMe*moeA*moeA*moeT*moeT | 20 |
| N02A058 | moeT*moeT*moeT*moeCMe*moeA*dG*dG*dT*dT*dT*dT*dT*dCMe*dCMe*dA*moeT*moeCMe*moeG*moeCMe*moeA | 20 |
| N02A063 | moeT*moeCMe*moeA*moeG*moeG*dT*dT*dT*dT*dT*dCMe*dCMe*dA*dT*dCMe*dG*moeCMe*moeA*moeCMe*moeA*moeT | 21 |
| N02A064 | moeT*moeCMe*moeA*moeA*moeT*dT*dG*dCMe*dT*dG*dT*dA*dA*dG*dA*moeT*moeT*moeG*moeT*moeCMe | 20 |
| N02A066 | moeCMe*moeA*moeG*moeG*moeT*dT*dT*dT*dT*dCMe*dCMe*dA*dT*dCMe*dG*dCMe*moeA*moeCMe*moeA*moeT*moeT | 21 |
| N02A076 | moeA*moeA*moeA*moeT*moeG*dG*dT*dA*dA*dT*dT*dG*dCMe*dA*dA*dG*dA*moeT*moeCMe*moeT*moeA*moeCMe | 22 |
| N02A077 | moeT*moeG*moeT*moeCMe*moeCMe*dA*dT*dT*dG*dG*dG*dG*dA*dG*dCMe*dA*dT*moeG*moeA*moeG*moeG*moeG | 22 |
| N02A081 | moeT*moeT*moeT*moeCMe*moeA*dG*dG*dT*dT*dT*dT*dT*dCMe*dCMe*dA*dT*moeCMe*moeG*moeCMe*moeA*moeCMe | 21 |
| N02A084 | moeCMe*moeT*moeT*moeT*moeCMe*dA*dG*dG*dT*dT*dT*dT*dT*dCMe*dCMe*moeA*moeT*moeCMe*moeG*moeCMe | 20 |
| N02A085 | moeCMe*moeCMe*moeA*moeT*moeCMe*dA*dT*dG*dT*dA*dA*dA*dCMe*dA*dA*dT*dA*moeA*moeG*moeG*moeCMe*moeA | 22 |
| N02A096 | moeT*moeT*moeCMe*moeA*moeG*dG*dT*dT*dT*dT*dT*dCMe*dCMe*dA*dT*dCMe*dG*moeCMe*moeA*moeCMe*moeA*moeT | 22 |
| N02A106 | moeCMe*moeT*moeA*moeT*moeG*dCMe*dA*dA*dA*dT*dG*dG*dT*dA*dA*dT*dT*moeG*moeCMe*moeA*moeA*moeG | 22 |
| N02A109 | moeCMe*moeT*moeT*moeCMe*moeCMe*dA*dT*dCMe*dT*dG*dT*dG*dA*dA*dCMe*dG*dA*moeA*moeG*moeA*moeG*moeA | 22 |
| N02A114 | moeT*moeG*moeG*moeT*moeA*dA*dT*dT*dG*dCMe*dA*dA*dG*dA*dT*moeCMe*moeT*moeA*moeCMe*moeA | 20 |
| N02A120 | moeCMe*moeA*moeG*moeG*moeT*dT*dT*dT*dT*dCMe*dCMe*dA*dT*dCMe*dG*moeCMe*moeA*moeCMe*moeA*moeT | 20 |
| N02A131 | moeT*moeT*moeCMe*moeA*moeG*dG*dT*dT*dT*dT*dT*dCMe*dCMe*dA*moeT*moeCMe*moeG*moeCMe*moeA | 19 |
| N02A133 | moeCMe*moeCMe*moeT*moeCMe*moeA*dT*dT*dCMe*dCMe*dT*dT*dCMe*dA*dA*dA*dT*moeCMe*moeT*moeA*moeG*moeA | 21 |
| N02A145 | moeCMe*moeT*moeCMe*moeA*moeT*dA*dG*dA*dA*dCMe*dT*dT*dG*dCMe*dCMe*dA*moeG*moeCMe*moeA*moeA*moeA | 21 |
| N02A153 | moeT*moeG*moeG*moeT*moeA*dA*dT*dT*dG*dCMe*dA*dA*dG*dA*dT*dCMe*moeT*moeA*moeCMe*moeA*moeA | 21 |
| N02A155 | moeG*moeCMe*moeT*moeG*moeT*dCMe*dCMe*dA*dT*dT*dG*dG*dG*dG*dA*dG*dCMe*moeA*moeT*moeG*moeA*moeG | 22 |
| N02A157 | moeCMe*moeT*moeT*moeT*moeCMe*dA*dG*dG*dT*dT*dT*dT*dT*dCMe*dCMe*dA*moeT*moeCMe*moeG*moeCMe*moeA | 21 |
| N02A158 | moeT*moeA*moeG*moeT*moeCMe*dCMe*dA*dA*dT*dT*dA*dG*dT*dG*dCMe*dA*dA*moeA*moeCMe*moeA*moeCMe*moeA | 22 |
| N02A168 | moeT*moeA*moeCMe*moeA*moeT*dA*dCMe*dCMe*dCMe*dT*dG*dA*dA*dT*dCMe*dT*moeG*moeT*moeG*moeCMe*moeT | 21 |
| N02A182 | moeCMe*moeT*moeG*moeT*moeCMe*dCMe*dA*dT*dT*dG*dG*dG*dG*dA*dG*dCMe*moeA*moeT*moeG*moeA*moeG | 21 |
| N02A186 | moeA*moeT*moeCMe*moeCMe*moeA*dA*dT*dA*dT*dG*dG*dA*dG*dA*dG*dCMe*dA*moeA*moeT*moeT*moeCMe*moeCMe | 22 |
| N02A191 | moeT*moeT*moeT*moeCMe*moeA*dG*dG*dT*dT*dT*dT*dT*dCMe*dCMe*moeA*moeT*moeCMe*moeG*moeCMe | 19 |
| N02A193 | moeT*moeT*moeA*moeT*moeCMe*dCMe*dA*dA*dT*dA*dT*dG*dG*dA*dG*dA*moeG*moeCMe*moeA*moeA*moeT | 21 |
| N02A199 | moeA*moeCMe*moeT*moeA*moeT*dG*dCMe*dA*dA*dA*dT*dG*dG*dT*dA*dA*dT*moeT*moeG*moeCMe*moeA*moeA | 22 |
| N02A207 | moeT*moeCMe*moeCMe*moeA*moeT*dCMe*dT*dT*dT*dT*dA*dT*dG*dT*dA*dT*moeA*moeT*moeA*moeCMe*moeT | 21 |
| N02A211 | moeT*moeT*moeCMe*moeA*moeG*dG*dT*dT*dT*dT*dT*dCMe*dCMe*dA*dT*dCMe*moeG*moeCMe*moeA*moeCMe*moeA | 21 |
| N02A218 | moeT*moeCMe*moeA*moeG*moeG*dT*dT*dT*dT*dT*dCMe*dCMe*dA*dT*dCMe*moeG*moeCMe*moeA*moeCMe*moeA | 20 |
| N02A228 | moeA*moeT*moeT*moeT*moeT*dA*dT*dCMe*dCMe*dA*dA*dT*dA*dT*dG*dG*moeA*moeG*moeA*moeG*moeCMe | 22 |
| N02A266 | moeCMe*moeT*moeG*moeT*moeCMe*dCMe*dA*dT*dT*dG*dG*dG*dG*dA*dG*dCMe*dA*moeT*moeG*moeA*moeG*moeG | 22 |
| N02A274 | moeT*moeCMe*moeT*moeCMe*moeCMe*dA*dT*dCMe*dT*dT*dT*dA*dT*dG*dT*dA*moeT*moeA*moeT*moeA*moeCMe | 21 |
| N02A277 | moeT*moeT*moeT*moeCMe*moeA*dG*dG*dT*dT*dT*dT*dT*dCMe*dCMe*dA*dT*dCMe*moeG*moeCMe*moeA*moeCMe*moeA | 22 |

| | | |
|---|---|---|
| N02A301 | moeT*moeA*moeG*moeT*moeCMe*dCMe*dA*dA*dT*dT*dA*dG*dT*dG*dCMe*dA*moeA*moeA*moeCMe*moeA*moeCMe | 20 |
| N02A315 | moeCMe*moeA*moeA*moeA*moeT*dG*dG*dT*dA*dA*dT*dT*dG*dCMe*dA*moeA*moeG*moeA*moeT*moeCMe | 22 |

d indicate DNA; * indicate phosphorothioate (PS) linkage; Me indicate 5-Methyl pyrimidine; moe indicate 2'-*O*-methoxyethyl (2'-MOE);
