## Supplementary material for "Sustained Relief of Chronic Pain via a Nav1.7-Targeting ASO–siRNA Conjugate": Supplental Table2

**Table S2: The sequences and modifications of siRNAs**

| siRNA # | Strand | Sequence(5' to 3') | Target exon |
|---|---|---|---|
| N02S101 | S | a*g*uccuCuAAGaagaauaucu | 3 |
| | AS | VPa*G*auaUuCUucuuAgAggacu*g*a | |
| N02S102 | S | g*u*ccucUaAGAagaauaucua | 3 |
| | AS | VPu*A*gauAuUCuucuUaGaggac*u*g | |
| N02S103 | S | u*c*cucuAaGAAgaauaucuau | 3 |
| | AS | VPa*U*agaUaUUcuucUuAgagga*c*u | |
| N02S104 | S | c*c*ucuaAgAAGaauaucuauu | 3 |
| | AS | VPa*A*uagAuAUucuuCuUagagg*a*c | |
| N02S105 | S | c*u*cuaaGaAGAauaucuauua | 3 |
| | AS | VPu*A*auaGaUAuucuUcUuagag*g*a | |
| N02S106 | S | u*c*uaagAaGAAuaucuauuaa | 3 |
| | AS | VPu*U*aauAgAUauucUuCuuaga*g*g | |
| N02S107 | S | u*a*agaaGaAUAucuauuaaga | 3 |
| | AS | VPu*C*uuaAuAGauauUcUucuua*g*a | |
| N02S108 | S | a*a*gaagAaUAUcuauuaagau | 3 |
| | AS | VPa*U*cuuAaUAgauaUuCuucuu*a*g | |
| N02S109 | S | a*u*uguuUuUGCguauuuaaca | 5-6 |
| | AS | VPu*G*uuaAaUAcgcaAaAacaau*g*a | |
| N02S110 | S | u*g*uuuuUgCGUauuuaacaga | 5-6 |
| | AS | VPu*C*uguUaAAuacgCaAaaaca*a*u | |
| N02S111 | S | g*u*uuuuGcGUAuuuaacagaa | 5-6 |
| | AS | VPu*U*cugUuAAauacGcAaaaac*a*a | |
| N02S112 | S | u*u*uuugCgUAUuuaacagaau | 5-6 |
| | AS | VPa*U*ucuGuUAaauaCgCaaaaa*c*a | |
| N02S113 | S | u*u*uugcGuAUUuaacagaauu | 5-6 |
| | AS | VPa*A*uucUgUUaaauAcGcaaaa*a*c | |
| N02S114 | S | u*u*uaacAgAAUuuguaaaccu | 6 |
| | AS | VPa*G*guuUaCAaauuCuGuuaaa*u*a | |
| N02S115 | S | u*u*aacaGaAUUuguaaaccua | 6 |
| | AS | VPu*A*gguUuACaaauUcUguuaa*a*u | |
| N02S116 | S | u*g*uguuUgCACuaauuggacu | 7 |
| | AS | VPa*G*uccAaUUagugCaAacaca*c*u | |
| N02S117 | S | g*u*guuuGcACUaauuggacua | 7 |
| | AS | VPu*A*gucCaAUuaguGcAaacac*a*c | |
| N02S118 | S | g*u*uugcAcUAAuuggacuaca | 7 |
| | AS | VPu*G*uagUcCAauuaGuGcaaac*a*c | |
| N02S119 | S | u*g*cacuAaUUGgacuacagcu | 7 |
| | AS | VPa*G*cugUaGUccaaUuAgugca*a*a | |
| N02S120 | S | c*a*cuaaUuGGAcuacagcugu | 7 |
| | AS | VPa*C*agcUgUAguccAaUuagug*c*a | |
| N02S121 | S | a*c*uaauUgGACuacagcuguu | 7 |
| | AS | VPa*A*cagCuGUagucCaAuuagu*g*c | |
| N02S122 | S | g*c*ccucAuGCUccccaaugga | 12 |
| | AS | VPu*C*cauUgGGgagcAuGagggc*u*g | |
| N02S123 | S | c*c*ucauGcUCCccaauggaca | 12 |

| | | | |
|---|---|---|---|
| | AS | VPu*G*uccAuUGgggaGcAugagg*g*c | |
| N02S124 | S | c*a*ugcuCcCCAauggacagcu | 12 |
| | AS | VPa*G*cugUcCAuuggGgAgcaug*a*g | |
| N02S125 | S | a*u*gcucCcCAAuggacagcuu | 12 |
| | AS | VPa*A*gcuGuCCauugGgGagcau*g*a | |
| N02S126 | S | g*c*ucccCaAUGgacagcuucu | 12 |
| | AS | VPa*G*aagCuGUccauUgGgagc*a*u | |
| N02S127 | S | g*g*aauuGcUCUccauauugga | 14 |
| | AS | VPu*C*caaUaUGgagaGcAauucc*a*g | |
| N02S128 | S | g*a*auugCuCUCcauauuggau | 14 |
| | AS | VPa*U*ccaAuAUggagAgCaauuc*c*a | |
| N02S129 | S | a*a*uugcUcUCCauauuggaua | 14 |
| | AS | VPu*A*uccAaUAuggaGaGcaauu*c*c | |
| N02S130 | S | a*u*ugcuCuCCAuauuggauaa | 14 |
| | AS | VPu*U*aucCaAUauggAgAgcaau*u*c | |
| N02S131 | S | u*u*gcucUcCAUauuggauaaa | 14 |
| | AS | VPu*U*uauCcAAuaugGaGagcaa*u*u | |
| N02S132 | S | u*g*cucuCcAUAuuggauaaaa | 14 |
| | AS | VPu*U*uuaUcCAauauGgAgagca*a*u | |
| N02S133 | S | g*c*ucucCaUAUuggauaaaau | 14 |
| | AS | VPa*U*uuuAuCCaauaUgGagagc*a*a | |
| N02S134 | S | c*u*cuccAuAUUggauaaaauu | 14 |
| | AS | VPa*A*uuuUaUCcaauAuGgagag*c*a | |
| N02S135 | S | c*u*ccauAuUGGauaaaauuca | 14 |
| | AS | VPu*G*aauUuUAuccaAuAuggag*a*g | |
| N02S136 | S | u*c*cauaUuGGAuaaaauucaa | 14 |
| | AS | VPu*U*gaaUuUUauccAaUaugga*g*a | |
| N02S137 | S | c*c*auauUgGAUaaaauucaaa | 14 |
| | AS | VPu*U*ugaAuUUuaucCaAuaugg*a*g | |
| N02S138 | S | g*a*uccuUuUGUagaucuugca | 14 |
| | AS | VPu*G*caaGaUCuacaAaAggauc*c*a | |
| N02S139 | S | a*u*ccuuUuGUAgaucuugcaa | 14 |
| | AS | VPu*U*gcaAgAUcuacAaAaggau*c*c | |
| N02S140 | S | u*c*cuuuUgUAGaucuugcaau | 14 |
| | AS | VPa*U*ugcAaGAucuaCaAaagga*u*c | |
| N02S141 | S | c*c*uuuuGuAGAucuugcaauu | 14 |
| | AS | VPa*A*uugCaAGaucuAcAaaagg*a*u | |
| N02S142 | S | c*u*uuugUaGAUcuugcaauua | 14 |
| | AS | VPu*A*auuGcAAgaucUaCaaaag*g*a | |
| N02S143 | S | u*u*guagAuCUUgcaauuacca | 14 |
| | AS | VPu*G*guaAuUGcaagAuCuacaa*a*a | |
| N02S144 | S | u*g*uagaUcUUGcaauuaccau | 14 |
| | AS | VPa*U*gguAaUUgcaaGaUcuaca*a*a | |
| N02S145 | S | g*u*agauCuUGCaauuaccauu | 14 |
| | AS | VPa*A*uggUaAUugcaAgAucuac*a*a | |
| N02S146 | S | u*a*gaucUuGCAauuaccauuu | 14 |
| | AS | VPa*A*augGuAAuugcAaGaucua*c*a | |

| | | | |
|---|---|---|---|
| N02S147 | S | a*u*cuugCaAUUaccauuugca | 14 |
| | AS | VPu*G*caaAuGGuaauUgCaagau*c*u | |
| N02S148 | S | u*c*uugcAaUUAccauuugcau | 14 |
| | AS | VPa*U*gcaAaUGguaaUuGcaaga*u*c | |
| N02S149 | S | c*u*ugcaAuUACcauuugcaua | 14 |
| | AS | VPu*A*ugcAaAUgguaAuUgcaag*a*u | |
| N02S150 | S | u*g*caauUaCCAuuugcauagu | 14 |
| | AS | VPa*C*uauGcAAauggUaAuugca*a*g | |
| N02S151 | S | g*c*aauuAcCAUuugcauaguu | 14 |
| | AS | VPa*A*cuaUgCAaaugGuAauugc*a*a | |
| N02S152 | S | a*u*uaccAuUUGcauaguuuua | 14 |
| | AS | VPu*A*aaaCuAUgcaaAuGguaau*u*g | |
| N02S153 | S | u*u*accaUuUGCauaguuuuaa | 14 |
| | AS | VPu*U*aaaAcUAugcaAaUgguaa*u*u | |
| N02S154 | S | u*a*ccauUuGCAuaguuuuaaa | 14 |
| | AS | VPu*U*uaaAaCUaugcAaAuggua*a*u | |
| N02S155 | S | g*u*gccuUaUUGuuuacaugau | 16 |
| | AS | VPa*U*cauGuAAacaaUaAggcac*a*u | |
| N02S156 | S | a*a*agacCaUUAagauuauccu | 20 |
| | AS | VPa*G*gauAaUCuuaaUgGucuuu*u*u | |
| N02S157 | S | g*a*ccauUaAGAuuauccugga | 20 |
| | AS | VPu*C*cagGaUAaucuUaAugguc*u*u | |
| N02S158 | S | c*c*auuaAgAUUauccuggagu | 20 |
| | AS | VPa*C*uccAgGAuaauCuUaaugg*u*c | |
| N02S159 | S | c*a*uuaaGaUUAuccuggagua | 20 |
| | AS | VPu*A*cucCaGGauaaUcUuaaug*g*u | |
| N02S160 | S | a*u*uaagAuUAUccuggaguau | 20 |
| | AS | VPa*U*acuCcAGgauaAuCuuaau*g*g | |
| N02S161 | S | u*a*gauuUgAAGgaaugagggu | 21-22 |
| | AS | VPa*C*ccuCaUUccuuCaAaucua*g*a | |
| N02S162 | S | u*u*gcugGcAAGuucuaugagu | 22 |
| | AS | VPa*C*ucaUaGAacuuGcCagcaa*a*c | |
| N02S163 | S | g*c*uggcAaGUUcuaugagugu | 22 |
| | AS | VPa*C*acuCaUAgaacUuGccagc*a*a | |
| N02S164 | S | a*u*gugcGaUGGaaaaaccuga | 22 |
| | AS | VPu*C*aggUuUUuccaUcGcacau*u*u | |
| N02S165 | S | u*g*ugcgAuGGAaaaaccugaa | 22 |
| | AS | VPu*U*cagGuUUuuccAuCgcaca*u*u | |
| N02S166 | S | g*u*gcgaUgGAAaaaccugaaa | 22 |
| | AS | VPu*U*ucaGgUUuuucCaUcgcac*a*u | |
| N02S167 | S | g*c*gaugGaAAAaccugaaagu | 22 |
| | AS | VPa*C*uuuCaGGuuuuUcCaucgc*a*c | |
| N02S168 | S | c*a*ugguAaCCAugaugguaga | 26 |
| | AS | VPu*C*uacCaUCauggUuAccaug*u*u | |
| N02S169 | S | a*u*gguaAcCAUgaugguagaa | 26 |
| | AS | VPu*U*cuaCcAUcaugGuUaccau*g*u | |
| N02S170 | S | g*g*uaacCaUGAugguagaaaa | 26 |

| | | | |
|---|---|---|---|
| | AS | VPu*U*uucUaCCaucaUgGuuacc*a*u | |
| N02S171 | S | g*a*uggaUuCUCuucguucaca | 27 |
| | AS | VPu*G*ugaAcGAagagAaUccauc*u*c | |
| N02S172 | S | u*g*gauuCuCUUcguucacaga | 27 |
| | AS | VPu*C*uguGaACgaagAgAaucca*u*c | |
| N02S173 | S | g*g*auucUcUUCguucacagau | 27 |
| | AS | VPa*U*cugUgAAcgaaGaGaaucc*a*u | |
| N02S174 | S | u*u*cucuUcGUUcacagaugga | 27 |
| | AS | VPu*C*cauCuGUgaacGaAgagaa*u*c | |
| N02S175 | S | u*c*ucuuCgUUCacagauggaa | 27 |
| | AS | VPu*U*ccaUcUGugaaCgAagaga*a*u | |
| N02S176 | S | u*c*uucgUuCACagauggaaga | 27 |
| | AS | VPu*C*uucCaUCugugAaCgaaga*g*a | |
| N02S177 | S | c*u*ucguUcACAgauggaagaa | 27 |
| | AS | VPu*U*cuuCcAUcuguGaAcgaag*a*g | |
| N02S178 | S | g*u*ucacAgAUGgaagaaaggu | 27 |
| | AS | VPa*C*cuuUcUUccauCuGugaac*g*a | |
| N02S179 | S | u*u*cacaGaUGGaagaaagguu | 27 |
| | AS | VPa*A*ccuUuCUuccaUcUgugaa*c*g | |
| N02S180 | S | c*a*cagaUgGAAgaaagguuca | 27 |
| | AS | VPu*G*aacCuUUcuucCaUcugug*a*a | |
| N02S181 | S | a*c*agauGgAAGaaagguucau | 27 |
| | AS | VPa*U*gaaCcUUucuuCcAucugu*g*a | |
| N02S182 | S | u*a*uauaCaUAAaagauggaga | 27 |
| | AS | VPu*C*uccAuCUuuuaUgUauaua*c*u | |
| N02S183 | S | u*a*uacaUaAAAgauggagaca | 27 |
| | AS | VPu*G*ucuCcAUcuuuUaUguaua*u*a | |

2′-*O*-C16

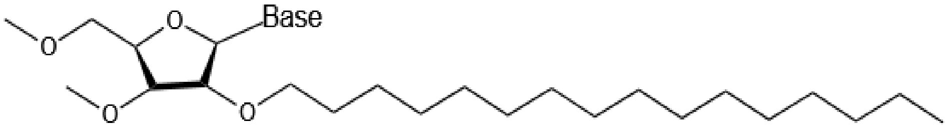


5'-(*E*)-VP

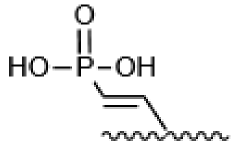


S and AS indicate sense and antisense strands, respectively; upper-case and lower-case letters indicate 2′-deoxy-2′-fluoro (2′-F) and 2′-*O*-methyl (2′-OMe) ribosugar modifications, respectively; * indicate phosphorothioate (PS) linkage; C16 indicate 2′-*O*-C16 ligand conjugated nucleotide (structure above); VP indicate 5'-(*E*)-vinylphosphonate (structure above)
