## Supplementary material for "Sustained Relief of Chronic Pain via a Nav1.7-Targeting ASO–siRNA Conjugate": Supplental Table3

**Table S3: The sequences and modifications of ASCs**

| siRNA # | Strand | Sequence(5'-3') |
| --- | --- | --- |
| N02C0701 | S | moeT*moeG*moeG*moeT*moeA*dA*dT*dT*dG*dCMe*dA*dA*dG*dA*dT*moeCMe*moeT*moeA*moeCMe*moeA*PEG6*mU*mA*mCmCmAmUfUmUfGfCfAmUmAmGmUmUmUmUmAmAmA |
| | AS | VPmU*fU*mUmAmAfAmAfCfUmAmUmGmCfAmAfAmUmGmGmUmA*mA*mU |
| N02C0702 | S | mU*mA*mCmCmAmUfUmUfGfCfAmUmAmGmUmUmUmUmAmAmA*PEG6*moeT*moeG*moeG*moeT*moeA*dA*dT*dT*dG*dCMe*dA*dA*dG*dA*dT*moeCMe*moeT*moeA*moeCMe*moeA |
| | AS | VPmU*fU*mUmAmAfAmAfCfUmAmUmGmCfAmAfAmUmGmGmUmA*mA*mU |
| N02C0703 | S | moeT*moeG*moeG*moeT*moeA*dA*dT*dT*dG*dCMe*dA*dA*dG*dA*dT*moeCMe*moeT*moeA*moeCMe*moeA*PEG6*mU*mA*mCmCmAmUfUmUfGfCfAmUmAmGmUmUmUmUmAmAmA*PEG6*moeT*moeG*moeG*moeT*moeA*dA*dT*dT*dG*dCMe*dA*dA*dG*dA*dT*moeCMe*moeT*moeA*moeCMe*moeA |
| | AS | VPmU*fU*mUmAmAfAmAfCfUmAmUmGmCfAmAfAmUmGmGmUmA*mA*mU |
| N02C0501 | S | moeCMe*moeT*moeT*moeT*moeCMe*dA*dG*dG*dT*dT*dT*dT*dT*dCMe*dCMe*dA*moeT*moeCMe*moeG*moeCMe*moeA*PEG6*mU*mA*mCmCmAmUfUmUfGfCfAmUmAmGmUmUmUmUmAmAmA |
| | AS | VPmU*fU*mUmAmAfAmAfCfUmAmUmGmCfAmAfAmUmGmGmUmA*mA*mU |
| N02C0502 | S | mU*mA*mCmCmAmUfUmUfGfCfAmUmAmGmUmUmUmUmAmAmA*PEG6*moeCMe*moeT*moeT*moeT*moeCMe*dA*dG*dG*dT*dT*dT*dT*dT*dCMe*dCMe*dA*moeT*moeCMe*moeG*moeCMe*moeA |
| | AS | VPmU*fU*mUmAmAfAmAfCfUmAmUmGmCfAmAfAmUmGmGmUmA*mA*mU |
| N02C0503 | S | moeCMe*moeT*moeT*moeT*moeCMe*dA*dG*dG*dT*dT*dT*dT*dT*dCMe*dCMe*dA*moeT*moeCMe*moeG*moeCMe*moeA*PEG6*mU*mA*mCmCmAmUfUmUfGfCfAmUmAmGmUmUmUmUmAmAmA*PEG6*moeCMe*moeT*moeT*moeT*moeCMe*dA*dG*dG*dT*dT*dT*dT*dT*dCMe*dCMe*dA*moeT*moeCMe*moeG*moeCMe*moeA |
| | AS | VPmU*fU*mUmAmAfAmAfCfUmAmUmGmCfAmAfAmUmGmGmUmA*mA*mU |

| | | |
|---|---|---|
| N02C1201 | S | moeCMe*moeT*moeT*moeT*moeCMe*dA*dG*dG*dT*dT*dT*dT*dT*dCMe*dCMe*dA*moeT*moeCMe*moeG*moeCMe*moeA*PEG6*mG*mC*mUmCmUmCfCmAfUfAfUmUmGmGmAmUmAmAmAmAmU |
| | AS | VPmA*fU*mUmUmUfAmUfCfCmAmAmUmAfUmGfGmAmGmAmGmC*mA*mA |
| N02C1202 | S | mG*mC*mUmCmUmCfCmAfUfAfUmUmGmGmAmUmAmAmAmAmU*PEG6*moeCMe*moeT*moeT*moeT*moeCMe*dA*dG*dG*dT*dT*dT*dT*dT*dCMe*dCMe*dA*moeT*moeCMe*moeG*moeCMe*moeA |
| | AS | VPmA*fU*mUmUmUfAmUfCfCmAmAmUmAfUmGfGmAmGmAmGmC*mA*mA |
| N02C1203 | S | moeCMe*moeT*moeT*moeT*moeCMe*dA*dG*dG*dT*dT*dT*dT*dT*dCMe*dCMe*dA*moeT*moeCMe*moeG*moeCMe*moeA*PEG6*mG*mC*mUmCmUmCfCmAfUfAfUmUmGmGmAmUmAmAmAmAmU*PEG6*moeCMe*moeT*moeT*moeT*moeCMe*dA*dG*dG*dT*dT*dT*dT*dT*dCMe*dCMe*dA*moeT*moeCMe*moeG*moeCMe*moeA |
| | AS | VPa*U*uuuAuCCaauaUgGagagc*a*a |
| N02C1401 | S | moeT*moeG*moeG*moeT*moeA*dA*dT*dT*dG*dCMe*dA*dA*dG*dA*dT*moeCMe*moeT*moeA*moeCMe*moeA*PEG6*mG*mC*mUmCmUmCfCmAfUfAfUmUmGmGmAmUmAmAmAmAmU |
| | AS | VPmA*fU*mUmUmUfAmUfCfCmAmAmUmAfUmGfGmAmGmAmGmC*mA*mA |
| N02C1402 | S | mG*mC*mUmCmUmCfCmAfUfAfUmUmGmGmAmUmAmAmAmAmU*PEG6*moeT*moeG*moeG*moeT*moeA*dA*dT*dT*dG*dCMe*dA*dA*dG*dA*dT*moeCMe*moeT*moeA*moeCMe*moeA |
| | AS | VPmA*fU*mUmUmUfAmUfCfCmAmAmUmAfUmGfGmAmGmAmGmC*mA*mA |
| N02C1403 | S | moeT*moeG*moeG*moeT*moeA*dA*dT*dT*dG*dCMe*dA*dA*dG*dA*dT*moeCMe*moeT*moeA*moeCMe*moeA*PEG6*mG*mC*mUmCmUmCfCmAfUfAfUmUmGmGmAmUmAmAmAmAmU*PEG6*moeT*moeG*moeG*moeT*moeA*dA*dT*dT*dG*dCMe*dA*dA*dG*dA*dT*moeCMe*moeT*moeA*moeCMe*moeA |
| | AS | VPmA*fU*mUmUmUfAmUfCfCmAmAmUmAfUmGfGmAmGmAmGmC*mA*mA |
