## Supplementary material for "Sustained Relief of Chronic Pain via a Nav1.7-Targeting ASO–siRNA Conjugate": Supplental Table4

**Table S4: Chemically modified controls, and primer sequences**

| siRNA | Sense (5'-3') | Antisense (5'-3') |
|---|---|---|
| siRNA NTC | mUmUmCmUmCmCfGmAfAfCfGmUmGmUmCmAmCmGmUdTdT | mAfCmGmUmGfAmCfAfCmGmUmUmCfGmGfAmGmAmAdTdT |

| ASO | Sequence (5'-3') |
|---|---|
| Tandem CpG | dT*dC*dG*dT*dC*dG*dT*dT*dT*dT*dG*dT*dC*dG*dT*dT*dT*dT*dG*dT*dC*dG*dT*dT |
| 558807 | cG*cCMe*cA*dT*dG*dT*dT*dCMe*dT*dCMe*dA*dCMe*dA*cT*cT*cA |
| ASO NTC | moeT*moeA*moeG*moeT*moeG*dCMe*dG*dG*dA*dCMe*dCMe*dT*dA*dC*dC*moeCMe*moeA*moeCMe*moeG*moeA |

| Gene name | Forward primer(5' - 3') | Reverse primer(5' - 3') | Probe sequence(5' - 3') | Species |
|---|---|---|---|---|
| *SCN9A* | *AAGTTTCCACCTTGGTGTCG* | *GATCCTATATCTCTTCCTCTG* | *ACCACTCAGCATTCGT* | Human |
| *GAPDH* | *TGCACCACCAACTGCTTAGC* | *GGCATGGACTGTGGTCATGAG* | *CTGGCCAAGGTCATCCATGACAACT* | Human |
| *Scn9a* | *GAACCAGGCCAACATCGAA* | *TACTCGTGAACTCAGCAGCG* | *AGAGTTAGAATTTCAGCAGATG* | Rat |
| *Gapdh* | *GGATGCAGGGATGATGTTC* | *TGCACCACCAACTGCTTAG* | *ATCACGCCACAGCTTTCCAGAGGG* | Rat |

| Gene name | Forward primer(5' - 3') | Reverse primer(5' - 3') | Species |
|---|---|---|---|
| *HSD11B1* | *AGTCAACTTCCTCAGTTACGTGGTC* | *GGAGCTGCTTGCATATGGACTAT* | Human |
| *TMEM92-AS1* | *CATCTCAGCTGGAAGGTGAGG* | *TGTATGCTTTGGCACTGTGTGTT* | Human |
| *PPIL6* | *CCTCTGCACTTTATGACGCACTCA* | *GTTTTGGGACACACATCACAGTATAGC* | Human |
| *KLHDC9* | *GAGTCATGGGAAAATTAAGGAGGAAC* | *ACAAAGGAGGAGATGTGCGAGTATC* | Human |
| *HID1* | *GCTGAACAAACCCTACTCAATCCG* | *GGCAGAGAAGAGGAACCAGGTG* | Human |
| *INHBE* | *CATCAGCTTTGCTACTGTCACAGACT* | *ATGTGGTGCTCAGCCAGGAG* | Human |
| *CCNA1* | *CCACCAACCAGTTTCTCCTTCA* | *TGTTCACAGTATAGTTTGCCAGGC* | Human |
| *SLC39A10* | *GTGCTGGATTGACAGGAGGAA* | *GACCAACAGCTGTGCCTATGA* | Human |
| *SCN9A* | *AAGTTTCCACCTTGGTGTCG* | *GATCCTATATCTCTTCCTCTG* | Human |
| *Gapdh* | *TGCACCACCAACTGCTTAGC* | *GGCATGGACTGTGGTCATGAG* | Human |

f indicate 2’-deoxy-2'-fluoro (2'-F); m indicate 2'-*O*-methyl(2-OMe);  d indicate DNA; * indicate phosphorothioate (PS) linkage; Me indicate 5-Methyl pyrimidine; c indicate constrained ethyl bridged nucleic acid (cEt); moe indicate 2'-*O*-methoxyethyl (2'-MOE).
