## Supplementary material for "Sustained Relief of Chronic Pain via a Nav1.7-Targeting ASO–siRNA Conjugate": Supplental Table5

**Table S5: Target sequences in psiCHECK-2-SM-reporters**

| Oligonucleotide name | Sequence (5'-3') |
|---|---|
| N02S133-SM | ctcgagAATGATGGAGGTGGAGATAAAAAATGATGGAGGTGGAGATAAAAAATGATGGAGGTGGAGATAAAAgcggccgc |
| N02S138-SM | ctcgagTATGATGCACCAGGAATCTTGCTATGATGCACCAGGAATCTTGCTATGATGCACCAGGAATCTTGCgcggccgc |
| N02S139-SM | ctcgagTATGAGCTACCTGCTTCTTGCATATGAGCTACCTGCTTCTTGCATATGAGCTACCTGCTTCTTGCAgcggccgc |
| N02S140-SM | ctcgagATAGAAGCACCTCGACTTGCAAATAGAAGCACCTCGACTTGCAAATAGAAGCACCTCGACTTGCAAgcggccgc |
| N02S141-SM | ctcgagAATGAAGCACCAGGATTGCAATAATGAAGCACCAGGATTGCAATAATGAAGCACCAGGATTGCAATgcggccgc |
| N02S142-SM | ctcgagTATGAAGCACCTGGATGCAATTTATGAAGCACCTGGATGCAATTTATGAAGCACCTGGATGCAATTgcggccgc |
| N02S143-SM | ctcgagTAAGAAGCTCGAGCAAATTACCTAAGAAGCTCGAGCAAATTACCTAAGAAGCTCGAGCAAATTACCgcggccgc |
| N02S145-SM | ctcgagAATGAAGCACCAGGTTTACCATAATGAAGCACCAGGTTTACCATAATGAAGCACCAGGTTTACCATgcggccgc |
| N02S146-SM | ctcgagAAAGATGGACCAGGATACCATTAAAGATGGACCAGGATACCATTAAAGATGGACCAGGATACCATTgcggccgc |
| N02S147-SM | ctcgagTATGAAGCACCAGGACATTTGCTATGAAGCACCAGGACATTTGCTATGAAGCACCAGGACATTTGCgcggccgc |
| N02S151-SM | ctcgagAATGTTGCTGGTGGAGCATAGTAATGTTGCTGGTGGAGCATAGTAATGTTGCTGGTGGAGCATAGTgcggccgc |
| N02S154-SM | ctcgagTAAGATGCACCAGGTGTTTTAATAAGATGCACCAGGTGTTTTAATAAGATGCACCAGGTGTTTTAAgcggccgc |

Note: underline = complementary sequence with seed region of the siRNA antisense strand; lower case = sequence of restriction enzyme site
